# A *Phelipanche ramosa* KAI2 Protein Perceives enzymatically Strigolactones and Isothiocyanates

**DOI:** 10.1101/2020.06.09.136473

**Authors:** Alexandre de Saint Germain, Anse Jacobs, Guillaume Brun, Jean-Bernard Pouvreau, Lukas Braem, David Cornu, Guillaume Clavé, Emmanuelle Baudu, Vincent Steinmetz, Vincent Servajean, Susann Wicke, Kris Gevaert, Philippe Simier, Sofie Goormachtig, Philippe Delavault, François-Didier Boyer

## Abstract

*Phelipanche ramosa* is an obligate root-parasitic weed threatening major crops in central Europe. For its germination, it has to perceive various structurally diverging host-exuded signals, including isothiocyanates (ITCs) and strigolactones (SLs). However, the receptors involved are still uncharacterized. Here, we identified five putative SL receptors in *P. ramosa*, of which PrKAI2d3 is involved in seed germination stimulation. We established the high plasticity of PrKAI2d3, allowing interaction with different chemicals, including ITCs. The SL perception mechanism of PrKAI2d3 is similar to that of endogenous SLs in non-parasitic plants. We provide evidence that the PrKAI2d3 enzymatic activity confers hypersensitivity to SLs. Additionally, we demonstrated that methylbutenolide-OH binds PrKAI2d3 and stimulates *P. ramosa* germination with a bioactivity comparable to that of ITCs. This study highlights that *P. ramosa* has extended its signal perception system during evolution, a fact to be considered in the development of specific and efficient biocontrol methods.

## Introduction

Witchweeds (*Striga* spp.) and broomrapes (*Orobanche* and *Phelipanche* spp.) are obligate root-parasitic plants belonging to the Orobanchaceae family and together comprise the most threatening weeds of the major domesticated crops worldwide^1^. At maturity a single plant releases up to 100,000 microscopic seeds, resulting in severe soil pollution^2^. Seed germination of these obligate parasites requires the strict recognition of host-exuded germination stimulants. Broomrapes and witchweeds are all highly sensitive to strigolactones (SLs) secreted by plants into the rhizosphere at picomolar doses^3^. The structural core of SLs is a tricyclic lactone, referred to as the ABC part in canonical SLs or as a structural variety in non-canonical SLs, invariably connected to an α,β-unsaturated furanone moiety (D ring) via an enol-ether bridge (Supplementary Figure 1**a**)^3^. Some germination stimulants are exclusive to specific host-parasite interactions, as illustrated by the unique ability of *Phelipanche ramosa* to germinate upon sensing isothiocyanates (ITCs). These glucosinolate-breakdown products are exuded by rapeseed (*Brassica napiisp*)^2,4^, on which *P. ramosa* has adapted in a decade^1,2^. Broomrapes are increasingly problematic in both intensity and acreage in Europe, North Africa, and Asia, and they are expected to dramatically expand to new territories in the near future^5^. To date, several physical, cultural, chemical, and biological approaches have been explored to control root-parasitic weeds, but no method has been found completely satisfactory^6^.

Besides their involvement in germination, SLs have a strong stimulating activity on arbuscular mycorrhizal fungi (Glomeromycotina) by promoting mitochondrial metabolism and hyphal branching^7^, thereby mediating the establishment of the oldest mutualistic interaction of land plants^8^. In addition to their role in rhizosphere signaling, SLs act as hormones *in planta* with pervasive roles throughout the plant development^9^ and, as such, are perceived in vascular plants by the α/β-hydrolase DWARF14 (D14)^10,11^. Biochemical analyses with recombinant D14 proteins by means of the synthetic SL analog GR24 revealed that the SL signal transduction requires GR24 cleavage^11,12^. One of the cleavage products, the D ring, may remain covalently attached to the receptor^13,14^, thereby probably allowing the recruitment of partners for downstream processes^15,16^. The required SL cleavage for signal transduction is still under debate^15,16^.

In *Arabidopsis thaliana*, D14 belongs to a small gene family, including KARRIKIN INSENSITIVE2/HYPOSENSITIVE TO LIGHT (AtKAI2/AtHTL) that shares the α/β-hydrolase catalytic triad. However, AtKAI2 regulates AtD14-independent processes, such as seed germination, and preferentially perceives (−)-GR24 that mimics non-natural SLs, karrikins^17^, and supposedly a still unknown endogenous ligand^18^. Interestingly, the *KAI2* gene family has expanded during the evolution of obligate parasitic-plant genomes^19^. For example, *Phelipanche aegyptiaca* and *Striga hermonthica* possess five and eleven *KAI2/HTL* genes, respectively. Two *KAI2* paralogs play a role in SL response in *P. aegyptiaca*^19^ and six *S. hermonthica* KAI2/HTL proteins are hypersensitive to SLs^19,20^, with ShHTL7 exhibiting the same perception mechanism as D14^21^. The remarkable expansion of the *KAI2* gene family in Orobanchaceae along with the capacity of *P. ramosa* to perceive ITCs let us to assume that KAI2 proteins might also perceive other germination stimulants. Here, we characterize PrKAI2d3 as a *P. ramosa* SL receptor. We demonstrate that it is able to perceive natural SLs by an enzymatically dependent mechanism contributing to its hypersensitivity to SLs. In addition, we show that PrKAI2d3 perceives ITCs as well as a wide range of SL analogs that thus were not found bioactive for hormonal SL functions. Together, these results suggest that PrKAI2 proteins evolved as hypersensitive and plastic receptors, enabling the parasitic plant to detect various host exudate metabolites and contributing to its dramatic success.

## Results

### Identification and gene expression profile of PrKAI2 homologs

Iterative nucleotide and amino-acid BLAST analyses were carried out on the recently published transcriptome of *P. ramosa* rapeseed strain^22^, with known *KAI2* and *D14* sequences of non-parasitic and parasitic plants as queries^19,20^. Each newly identified sequence was used as a query for new BLAST searches to find all complete sequences. Duplicates were eliminated before alignment, allowing the detection of one *PrD14* and five *PrKAI2* putative orthologs that were amplified and re-sequenced from cDNA isolated from germinated seeds of the same *P. ramosa* strain. Maximum-likelihood analysis indicated that the predicted PrD14 and PrKAI2 proteins (Figure 1**a**) and genes (Supplementary Figure 2) belonged to clearly distinct D14 and KAI2 clades, respectively, with a conserved catalytic triad and an overall conserved environment of known active-site amino-acid residues (Supplementary Figure 3). Despite a poor separation of non-parasitic and parasitic KAI2 groups, the phylogenetic analysis strongly suggested that among the retrieved transcripts no PrKAI2 protein belonged to the intermediate KAI2i subclade, as previously described for other members of the *Orobanche* and *Phelipanche* genera^19^. The newly identified PrKAI2 and PrD14 proteins consistently clustered together with *P. aegyptiaca* sequences. Together with the reported low levels of genomic divergence in *Phelipanche*^23,24^, these findings hint at the detection of *P. aegyptiaca* protein orthologs. Hence, the *P. ramosa* sequences were renamed according to the current nomenclature^19^, namely one PrKAI2 protein belonging to the conserved KAI2 (PrKAI2c) clade and four PrKAI2 proteins from the divergent KAI2 clade (PrKAI2d1-d4).

**Figure 1.**
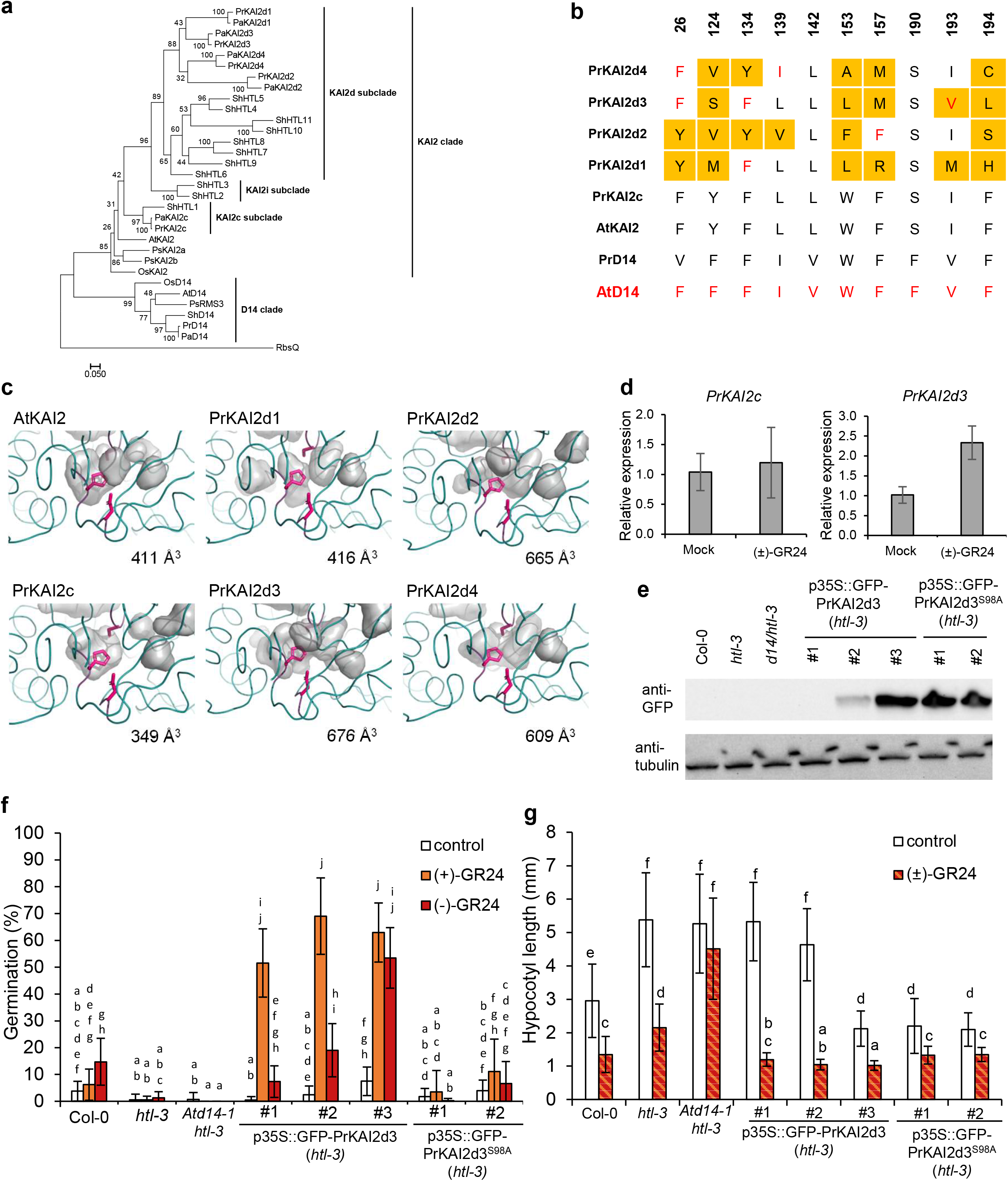
Identification of PrKAI2 putative SL receptors in *P. ramosa*. **(a)** Phylogenetic analysis of KAI2 and D14 amino acid sequences. The phylogenetic tree was constructed with the maximum likelihood method and 1,000 bootstraps replicates by means of RAxML. The scale bar represents 0.05 substitutions per site. Clades were designated as described^19^ **(b)** Amino acid sequence alignment of the active PrKAI2 protein sites. Amino acids that differed from AtKAI2 and those similar to AtD14 are in orange and red, respectively; whereas. AtHTL and PrKAI2d3 are in blue and yellow, respectively. A fully expanded alignment can be found in Supplementary Figure 3. **(c)** Visual representation of the ligand pockets of the *P. ramosa KAI2* genes. The KAI2 protein sequences were modeled with the chain A of the karrikin-bound *Arabidopsis* KAI2 structure as a template (PDB: 4JYP). The protein structures were generated with PYMOL and the cavities within the homology models were visualized with the surface mode on the setting “cavities and pockets culled” within PYMOL. **(d)** Expression of the *KAI2* genes in *P. ramosa*. Primers for *PrKAI2c* and *PrKAI2d3* were used in a qRT-PCR experiment with *EF101* as housekeeping gene on seeds treated for 1 h with (±)- GR24^76^ (*P* < 0.001, Student’s *t* test). **(e-g)** Cross-species complementation assays of the *htl-3* mutant with *P. ramosa* KAI2d3 and the catalytic site mutant S98A. **(e)** Protein levels of PrKAI2d3 of 4-day-old seedlings transformed with *p35S::GFP:PrKAI2d3* or *p35S::GFP:PrKAI2d3*^S98A^ detected with anti-GFP (top) and anti-tubulin (bottom) antibodies as loading control. The experiment was repeated twice with comparable results and one representative repeat is shown. (**f)** Seed germination after 5 days of growth at 25°C in the dark, with DMSO (control), 10 μM (+)-GR24, or 10 μM (−)-GR24 treatments. Transgenes were expressed in the null *htl-3* mutant background (Col-0 accession) under control of the 35S promoter. (one representative experiment of 18 wells per condition with an average of 19 seeds/well shown). **(g)** Hypocotyl lengths of 4-day-old seedlings grown under continuous red light at 21 °C (*n* = 25) with 10 μM (±)-GR24 treatments. Graphs represent means of three biological repeats ± SE. Statistical groups indicated by letters were determined by Kruskal-Wallis test with Dunn’s post test, *P* < 0.001 (**f)** and *P* < 0.05 (**g)**.

### PrKAI2d3 is a putative SL receptor

Sequence homology analysis of the ligand-binding pocket amino-acid residues of the PrKAI2 proteins with AtD14 and AtKAI2 showed that PrKAI2d3 possesses the highest similarity with AtD14 (Figure 1**b**). Predictive models generated *via* the SWISS-MODEL webserver revealed that the binding pocket of PrKAI2c is smaller than that of its putative ortholog AtKAI2 (Figure 1**c**) and that the binding pockets of the PrKAI2d receptors are wider than those of members of the conserved KAI2 clade, probably due to the absence of large hydrophobic residues (Figure 1**c**). Among all the divergent receptors PrKAI2d3 had the largest predicted binding pocket (Figure 1**c**). Finally, the expression level of *PrKAI2d3* transcripts in *P. ramosa* seeds was significantly up-regulated upon 1 hour of exogenous treatment with GR24, whereas the transcript levels for PrKAI2c, the putative AtKAI2 ortholog, did not change (Figure 1**d**). These properties are in agreement with a role of PrKAI2d3 as SL receptor.

As no mutants and nor an easy and fast transformation method are available for holoparasitic Orobanchaceae, it is difficult^25^ to validate the biological function of PrKAI2d3 directly in *P. ramosa*. Instead, we assayed the complementation of the *Arabidopsis htl-3* mutant phenotype with PrKAI2d3, a previously successful approach^19,20^. To assess the role of the catalytic triad in the perception process, *htl-3* was also transformed with a mutated PrKAI2d3^S98A^-encoding version. We used three p35S::GFP-PrKAI2d3 lines displaying various protein levels and two p35S::GFP-PrKAI2d3^S98A^ lines with a high protein expression (Figure 1**e**) to phenotype both germination and hypocotyl length.

In a thermo-inhibition assay^26^, almost no germination was detected for all lines upon control treatments. Germination of *Arabidopsis* (accession Columbia-0 [Col-0]) seeds was significantly stimulated upon 10 μM (−)-GR24, but not upon (+)-GR24, whereas none of the enantiomers had an effect on *htl-3* or *Atd14-1/htl-3* mutants (Figure 1**f**). Germination of all three p35S::GFP-PrKAI2d3 lines was strongly induced by (+)-GR24. Exogenous (−)-GR24 also significantly stimulated germination of these three lines (Figure 1**f**) and the response amplitude correlated positively with the protein abundance (Figure 1**e**). Analysis of a fourth p35S::GFP-PrKAI2d3 line showed that even a mild protein expression increased the *Arabidopsis* sensitivity a 100-fold towards (−)-GR24, enabling it to perceive up to picomolar doses of (+)-GR24 and, hence, to have the *P. ramosa* sensitivity (Figure 2**a-b**, Supplementary Figure 4, Supplementary Table 1). None of the two p35S::GFP-PrKAI2d3^S98A^ lines were sensitive to either (+)- or (−)-GR24.

**Figure 2.**
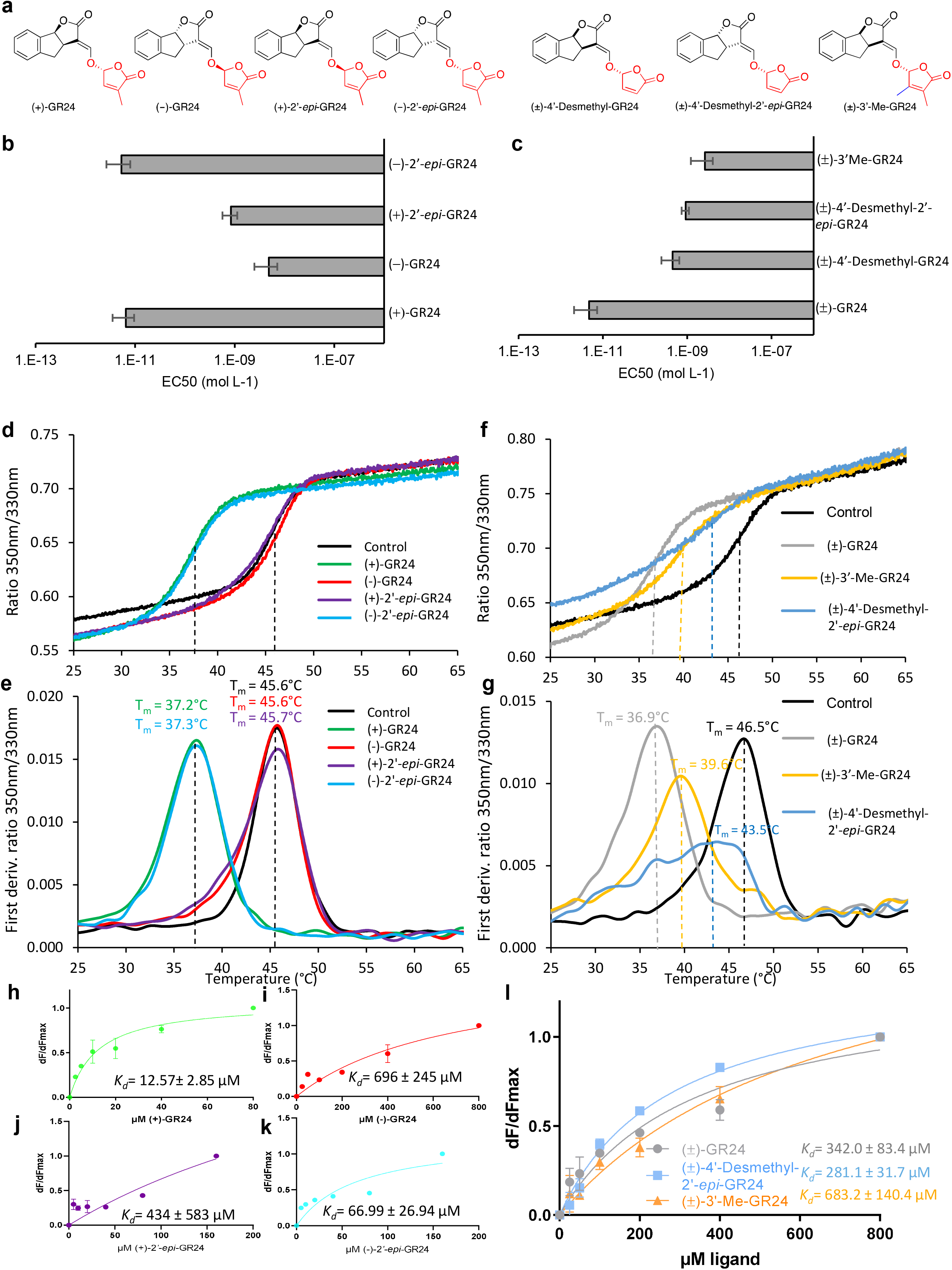
PrKAI2d3 shows a stereoselectivity towards GR24 analogs mimicking the SL natural stereoconfigurations and perceives SL analogs without a methyl group on the D-ring. **(a)** Structures of GR24 analogs. (**b)** and (**c**) Germination stimulation activity on *P. ramosa* seeds (EC_50_) with four stereoisomers and with GR24 analogs harboring variation on the D-ring (± SE), respectively. **(d-g)** Thermostability of PrKAI2d3 at 10 μM in the absence of ligands (black line) or in the presence of various ligands at 100 μM analyzed by nanoDSF. The panels **(d**) and (**f)** show the changes in fluorescence (ratio F_350nm_/F_330nm_) with temperature and **(e)** and (**g)** the first derivatives for the F_350nm_/F_330nm_ curve against the temperature gradient from which the apparent melting temperatures (Tm) was determined for each sample. The experiment was carried out twice. **(h)** and (**i)** SL analogs binding PrKAI2d3 based on intrinsic tryptophan fluorescence. Plots of fluorescence intensity *versus* probe concentrations. The change in intrinsic fluorescence was monitored (Supplementary Figure 8) and used to determine the apparent *K*D values. The plots represent the mean of two replicates and the experiments were repeated at least three times. The analysis was done with GraphPad Prism 5.0 Software.

Under red-light conditions, hypocotyls of *htl-3* and *Atd14-1*/*htl-3* mutants are more elongated than those of Col-0^17^ (Figure 1**g**). Application of (±)-GR24 significantly shortened the hypocotyls of Col-0 and the *htl-3* genotypes, but not those of the *Atd14-1/htl-3* plants. This observation corroborates a previous report that seedling photomorphogenesis is redundantly controlled by D14 and KAI2/HTL in *Arabidopsis*^17^. Interestingly, only the hypocotyls of the p35S::GFP-PrKAI2d3 and p35S::GFP-PrKAI2d3^S98A^ lines with the highest protein levels were shorter than those of the *htl-3* and *Atd14-1/htl-3* mutants under control conditions. However, the hypocotyls were significantly shorterned in all transformed lines after treatment with 1 M (±)-GR24 (Figure 1**g**). In summary, these data indicate that the PrKAI2d3 protein expression rescues the *htl-3* mutant phenotypes, with the catalytic Ser98 being essential for germination, but not for seedling photomorphogenesis.

### A structure-activity relationship study reveals that *P. ramosa* efficiently responds to SL analogs and mimics with various structures

As *P. ramosa* perceives many high structurally divergent germination stimulants, various receptors might underlie this plasticity^27,28^. To link the SL structural features with the *P. ramosa* seed germination activity, we carried out a structure-activity relationship (SAR) study through a rapid bioassay^29^.

First, we determined the *P. ramosa* sensitivity towards several GR24 analogs with varying stereogenic centers (Figure 2**a**). The lowest EC_50_ values were obtained with (+)-GR24 (6.5 pM) and (−)-2’-*epi*-GR24 (5.3 pM), of which the stereochemistry corresponds to natural canonical SLs of the strigol-type and orobanchol-type series, respectively (Supplementary Figure 1**a**). In contrast, *P. ramosa* was approximately 100-fold less sensitive to (−)-GR24 and (+)-2’-*epi*-GR24, of which the stereochemistry is not encountered in natural SLs (Figure 2**b**, Supplementary Figure 1**a**).

Second, we investigated whether substitutions occurring on the D-ring alter *P. ramosa* responses^27^. Surprisingly, both GR24 analogs without a methyl group on the D-ring, (±)-4’-desmethyl-2’-*epi*-GR24 and (±)-4’-desmethyl-GR24, significantly stimulated *P. ramosa* seed germination (EC_50_ = 0.92 and 0.45 nM), still with 100-fold lower EC_50_ values than those of (±)-GR24 (Figure 2**c**). Noteworthy, none of these substituted GR24 analogs could inhibit shoot branching even at 1 μM^27^. Inversely, *P. ramosa* was less sensitive to (±)-3’-methyl-GR24 (EC_50_ = 2.6 nM), which harbors two methyl groups on the D-ring (Figure 2**c**, Supplementary Figure 5, Supplementary Table 2) and is highly bioactive in repressing pea (*Pisum sativum*) shoot branching via RMS3/PsD14^13,27^.

Finally, the effect of the ABC-ring fragment substitution on the *P. ramosa* germination was analyzed. Profluorescent probes were used, in which the ABC-ring was replaced by a coumarine moiety (DiFMU) and one, two, or no methyl groups were added on the D-ring (GC240, GC242, and GC486, respectively)^13^. In another profluorescent probe, the ABC-ring was switched by the fluorescein moiety Yoshimulactone green (YLG)^30^ (Supplementary Figure 1**b**). Except for DiFMU, all profluorescent probes had a stimulating activity on the *P. ramosa* germination, but still with 1,000-to 10,000-fold lower EC_50_ values than those of (±)- GR24 (Supplementary Figure 6**a-c**, Supplementary Table 2).

Altogether, these results demonstrate that the stereochemistry of GR24 analogs is crucial for bioactivity and determines the sensitivity to germination-inducing substances. The sensitivity of *P. ramosa* towards desmethyl and 3’-methyl D-ring derivatives highlights differences in contrast to other vascular plants, particularly regarding the signaling via D14 for shoot branching. Additionally, the relative bioactivity of the profluorescent SL probes for germination stimulation in broomrape confirms that ABC-rings are not required for bioactivity^28^.

### SL analogs and mimics interact with PrKAI2d3 according to their germination stimulation activity

The questions arising from previous results are whether PrKAI2d3 has the ability to perceive such a large range of structurally divergent compounds and whether its interaction with these molecules is affected by mutations in the catalytic triad. To this end, we expressed and purified the PrKAI2d3 and PrKAI2d3^S98A^ proteins *in vitro* and assessed their abilities to interact with SLs and other chemical mediators.

Interactions between PrKAI2d3 and the SL analogs and profluorescent SL probes were analyzed with nano differential scanning fluorimetry (nanoDSF) by recording changes in the tryptophan fluorescence (ratio 350 nm/330 nm). In contrast to “classical” DSF^11^, nanoDSF does not require a dye and highlights interactions that do not induce conformational changes. Analysis of the initial fluorescence ratios revealed that the four GR24 stereoisomers interacted with PrKAI2d3 according to their bioactivity (Figure 2**d**). However, only (+)-GR24 and (−)- 2’-*epi*-GR24, the most bioactive analogs with natural configurations, induced a 8.5 °C decrease in the PrKAI2d3 melting temperature (Figure 2**e**), consistent with ligand-mediated protein destabilization.

When the GR24 analogs with varying methyl groups on their D-ring were used, all the analogs interacted and destabilized PrKAI2d3, although with an efficiency lower than that of (±)-GR24, and especially that of (±)-3’-Me-GR24 (Figure 2**f,g**). Similar shifts in melting temperatures of the PrKAI2d3 protein were observed with DSF, but without destabilization of the mutated PrKAI2d3^S98A^ protein (Supplementary Figure 7). The bioactive profluorescent probe GC242, used as a racemic mixture, or separate pure enantiomers also induced a shift in the PrKAI2d3 protein melting temperature (Supplementary Figure 6**d-i**). These results suggest that the enzymatic activity through Ser98 is required to destabilize the protein.

Next, we estimated the PrKAI2d3 affinity for GR24 analogs by means of the tryptophan intrinsic fluorescence assay. The affinity was higher for (+)-GR24 and (−)-2’-*epi*-GR24 (*K*_D_ = 12.57 ± 2.85 μM and 66.99 ± 26.94 μM, respectively) than for (−)-GR24 and (+)-2’-*epi*-GR24 (*K*_D_ = 696 ± 245 μM and 434 ± 593 μM, respectively), in accordance with their bioactivity on the *P. ramosa* germination (Figure 2**h-k**, Supplementary Figure 8), whereas the lower affinity of (±)-3’-Me-GR24 than that of (±)-4’-desmethyl-2’-*epi*-GR24 and (±)-GR24 is consistent with their reduced bioactivity range (Figure 2**l**). However, protein affinities were significantly lower (micromolar range) than the observed sensitivity of *P. ramosa* (picomolar range). As similar patterns had been reported for *Striga hermonthica*^30^, we tested this apparent contradiction by investigating the enzymatic activity of the PrKAI2d3 protein.

### The PrKAI2d3 enzymatic activity is associated with SL hypersensitivity

First, we tried to visualize the PrKAI2d3 hydrolase activity with a generic substrate, *para*-nitrophenyl acetate (*p*-NPA). Surprisingly, no hydrolytic activity was detected (Supplementary Figure 10) in contrast to other SL receptors with the same probe^13^. When PrKAI2d3 was incubated with (+)-GR24 and (−)-2’-*epi*-GR24, a cleavage activity was unambiguously observed by ultraperformance liquid chromatography (UHPLC)/UV DAD analysis (Figure 3**a**), but, opposite to AtD14 and AtKAI2, PrKAI2d3 did not cleave (−)-GR24 or (+)-2’-*epi*-GR24. Interestingly, a residual cleavage activity of PrKAI2d3^S98A^ still occurred with all GR24 isomers, but without stereoselectivity, possibly the reason for the partial complementation of *htl-3* with PrKAI2d3^S98A^-overproducing constructs (Figure 1**g**). These data suggest that SL hydrolysis and the subsequent signal transduction could happen without an intact catalytic triad.

**Figure 3.**
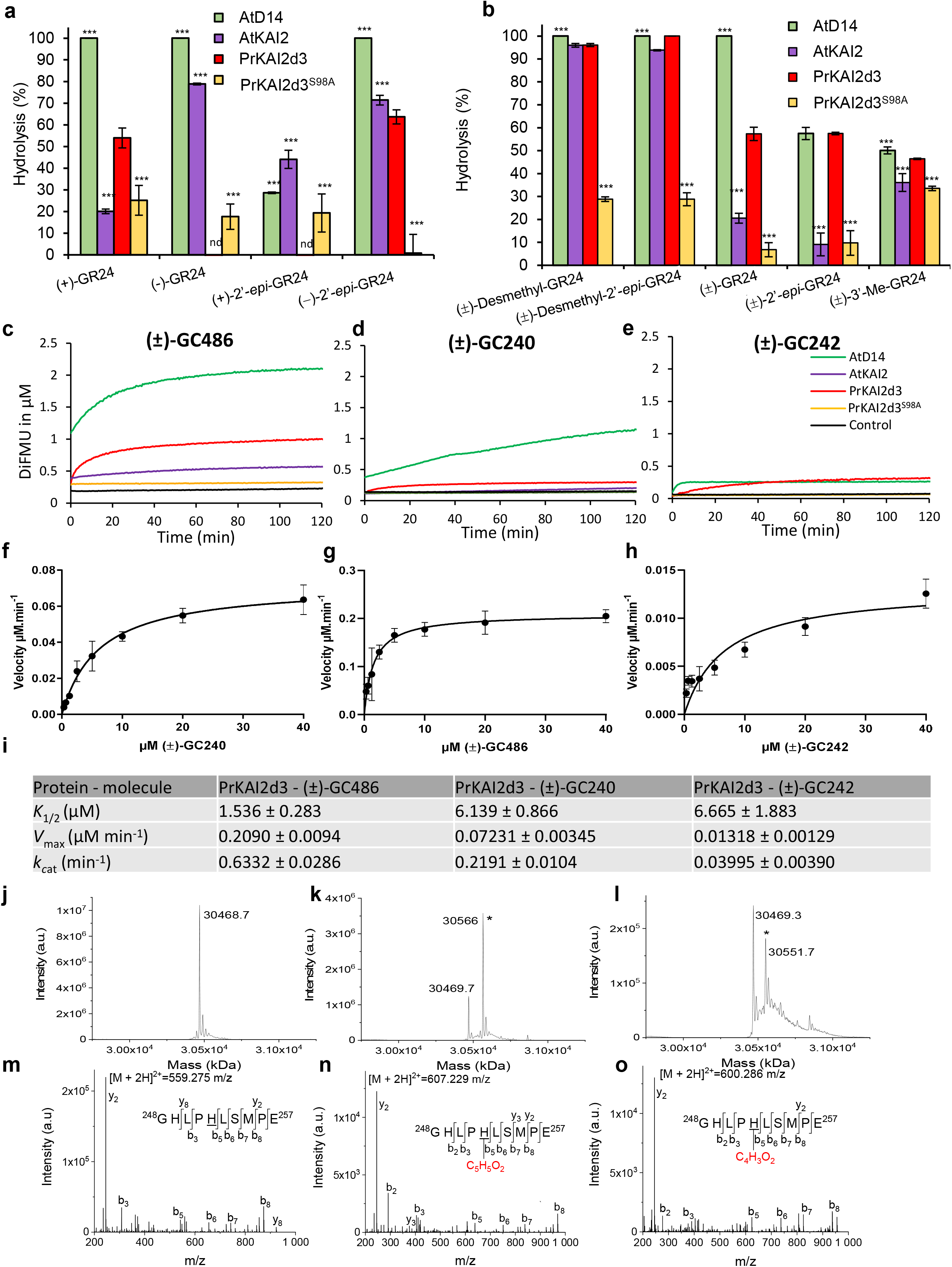
The PrKAId3 enzymatic activity improves the SL biological activity. (**a)** and (**b**) Hydrolysis activity of GR24 isomers and analogs by various proteins. (+)-GR24, (−)-GR24, (+)-2’*epi*-GR24 and (−)-2’*epi*-GR24 (**a**) and (±)-GR24, (±)-4’-desmethyl-GR24, (±)-4’-desmethyl-2’-*epi*-GR24, and (±)-3’-Me-GR24 (**b**) at 10 μM were incubated with PrKAI2d3, PrKAI2d3^S98A^, AtD14, and AtKAI2 at 5 μM for 150 min at 25 °C. UPLC-UV (260 nm) analysis was used to detect the remaining amount of GR24 isomers and analogs. Bars represent the mean value of the hydrolysis rate calculated from the remaining GR24 isomers and analogs taking into account the hydrolysis in the buffer alone (without protein sample), quantified with indanol as internal standard. Error bars represent the SD of three replicates (means ± SD, *n* = 3). nd, no hydrolysis detected. The asterisks indicate statistical significance from the PrKAI2d3 protein sample as \*\*\**P* ≤ 0.001; and *P* > 0.05, as measured by Kruskal-Wallis test. **(c-e)** Enzymatic kinetics for PrKAI2d3, PrKAI2d3^S98A^, AtD14 and AtKAI2 proteins incubated with (±)-GC486 **(c)**, (±)-GC240 **(d),** and (±)-GC242 **(e)**. Progress curves during hydrolysis of the probes, monitored (λe_m_ 460 nm) at 25 °C with the use of 400 nM protein and 20 μM probes. The traces represent one of the three replicates and the experiments were repeated at least twice. **(f-h)** Hyperbolic plot of the PrKAI2d3 presteady-state kinetics reaction velocity with (±)-GC486 **(f)**, (±)-GC240 **(g),** and (±)-GC242 **(h)**. The initial velocity was determined with profluorescent probe concentrations from 0.3 μM to 40 μM and with proteins at 400 nM. Error bars represent SE of the mean of three replicates and the experiments were repeated at least three times. **(i)** Kinetics constants of probes towards PrKAI2d3. *K*_1/2_ and *k*_cat_ are presteady-state kinetics constants for PrKAI2d3 with different profluorescent probes and represent the mean ± SE of three replicates. **(j-o)** Mass spectrometry characterization of covalent PrKAI2d3-ligand complexes. On the left, deconvoluted electrospray mass spectra of PrKAI2d3 prior and after addition of different ligands (±)-GR24 and (±)-GC486. Peaks with an asterisk correspond to PrKAI2d3 covalently bound to a ligand (PrKAI2d3-ligand). The mass increments were measured for different PrKAI2d3-ligand complexes: 96.3 Da (±)-GR24) and 82.4 Da (±)-GC486). Ligand-modified amino acids were identified by nanoLC-MSMS analyses after Glu-C proteolysis. On the right, fragmentation spectra of unmodified and different ligand-modified peptides. Labeled peaks correspond to the b and y fragments of the double-charged precursor ion displayed at the top. The histidine residue modified by different ligands is underlined.

As for the interaction assays, we tested the hydrolysis of analogs with substitutions on the D-ring. Desmethyl GR24 isomers were more efficiently hydrolyzed than GR24 by AtKAI2, PrKAI2d3, and PrKAId3^S98A^ proteins (Figure 3**b**). Additionally, these three purified proteins along with AtD14 also displayed a low, but significant, hydrolysis activity towards 3’-Me-GR24. These results indicate that PrKAI2d3 possesses an important hydrolysis capacity towards the natural configuration-mimicking GR24, albeit to a lesser extent than AtD14.

An enzymatic kinetic characterization of the purified proteins was carried out with the bioactive profluorescent probes as substrate. Monitoring the DiFMU fluorescence revealed that PrKAI2d3 hydrolyzed (±)-GC240, (±)-GC242, and (±)-GC486 (Figure 3**c-e**). For all probes, we observed a biphasic time course of fluorescence, consisting of a burst phase (also called initial phase), followed by a plateau phase (or slow phase for AtD14 incubated with (±)-GC240). In all cases, the plateau did not reach the maximum of the expected product (20 M), but a concentration product closer to the protein concentration (0.4 M) (Figure 3**c-e**), implying that PrKAI2d3 might act as a single turnover enzyme towards all GC probes tested, including GC486 that lacks the 3’-methyl group on the D-ring and has a Michaelian cleavage kinetic with AtD14 and RMS3/PsD14 proteins^13^. The S98A substitution in the catalytic triad drastically reduced the (±)-GC240 and (±)-GC242 cleavage, although the residual activity towards (±)-GC486 remained statistically significant. These observations confirmed that the mutation in the catalytic triad does not fully abolish the cleavage activity towards a compound without methyl on the D-ring.

Regarding single turnover enzymes, we defined *k_cat_* as the rate constant of the presteady-state phase (initial phase) and *K*_1/2_ as the probe concentration with half maximal velocity (*V_max_*) (Figure 3**g-i**). The similar *K*_1/2_ values of PrKAI2d3 with (±)-GC240 and (±)-GC242 (5.74 μM and 4.60 μM, respectively) confirmed the low influence of the C3’ methyl chain on the substrate affinity, but the differences in the *V_max_* (0.072 M.min^−1^ and 0.013 M.min^−1^, respectively) indicated that the C3’ methyl chain reduces the catalytic activity, corresponding with the bioactivities of these probes. Higher *K*1/2 values of PrKAI2d3 for (±)-GC240 than that for (±)-GC486 (5.74 μM and 1.53 μM, respectively) and *V_max_* values (0.072 M.min^−1^ and 0.209 M.min^−1^, respectively) highlighted the importance of the C4’ methyl chain in binding and catalytic affinity, supporting the results of the germination bioassays (Supplementary Figure 6**a-c**).

To test the hypothesis that PrKAI2d3 forms a stable intermediate with the D-ring, as previously demonstrated for other SL receptors (AtD14, PsD14/RMS3, ShHTL7, and D14)^13,14,21,31^, we incubated two bioactive ligands (±)-GR24 and probe (±)-GC486 with PrKAI2d3 at a pH of 6.8 and recorded mass spectrometry spectra under denaturing conditions. In all cases a mass shift occurred corresponding to the D-ring covalently bound to the protein (Figure 3**j-l**) and specifically attached to His249 of the catalytic triad (Figure 3**m-o**).

### ITCs interact with PrKAI2d3 and generate a covalent adduct to the catalytic serine

Although PrKAI2d3 perceives structurally diverging SL analogs and mimics that stimulate seed germination of *P. ramosa*, the specificity of the interaction between rapeseed and *P. ramosa* correlates with the parasite’s ability to perceive ITCs, which structurally differ greatly from SLs^4^. Therefore, we evaluated the ability of PrKAI2d3 to perceive ITCs. Seeds of *P. ramosa* were approximately 10,000-fold less sensitive to 2-PEITC and BITC than to (±)-GR24 (Figure 4**a-c**). Investigation of the putative interactions between PrKAI2d3 and ITCs by nanoDSF revealed a small shift (1.2-2.0 °C) in the PrKAI2d3 melting temperature upon high ITC concentrations, as further confirmed by “classical DSF” (Figure 4**d-e**, Supplementary Figure 11). These data indicated that ITCs interact with PrKAI2d3 with low affinity, corresponding with a stimulating potential on *P. ramosa* seeds lower than that of bioactive SLs^4^ and analogs. Finally, the apparent melting temperatures of PrKAI2d3 with BITC and 2-PEITC (44.3 °C and 45.1 °C, respectively) varied from values obtained with (±)-GR24 (36.9 °C), suggesting that ITCs might induce a conformational change that differs from the GR24-induced destabilization.

**Figure 4.**
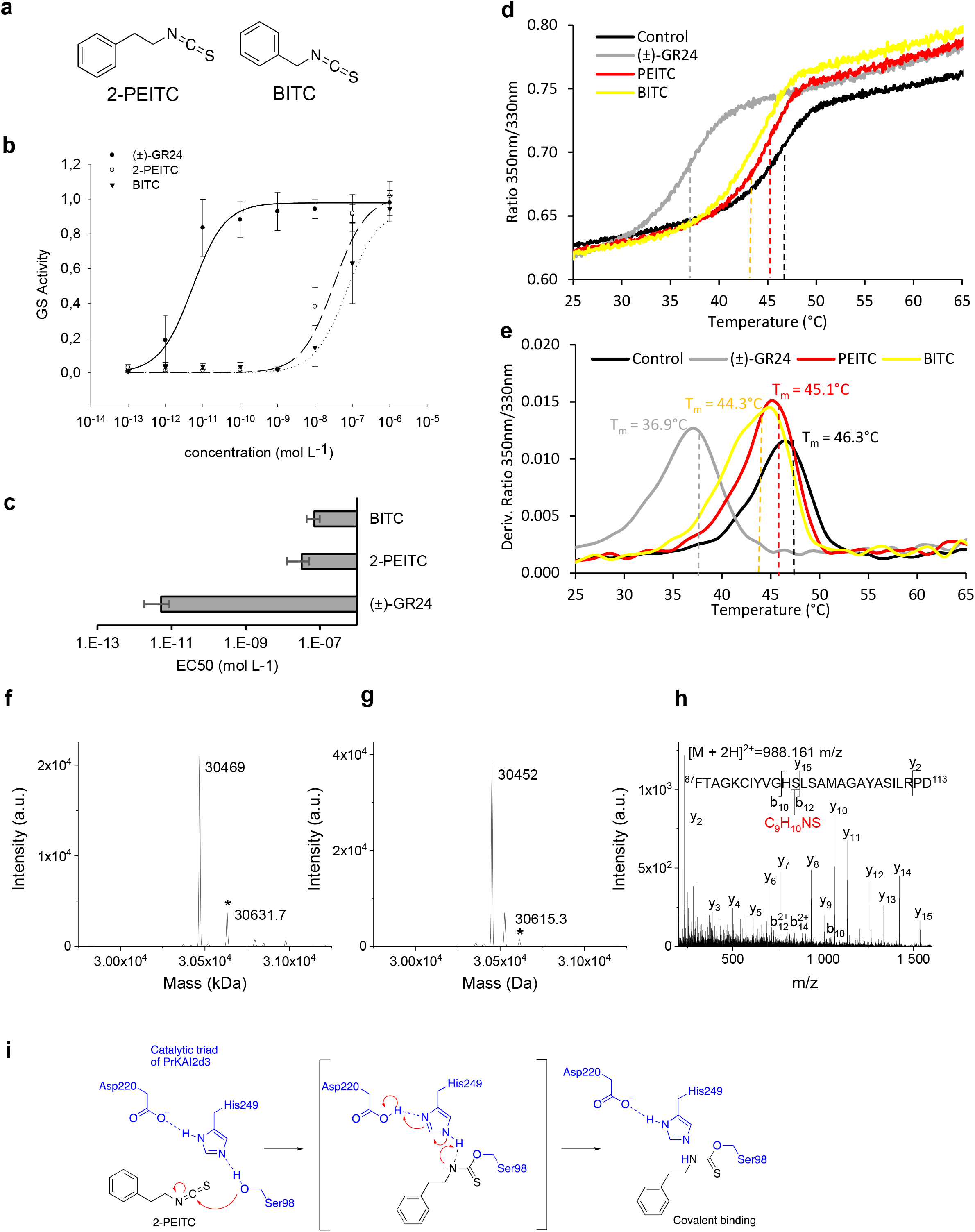
The isothiocyanate germination stimulants are perceived by PrKAI2d3. **(a)** Structure of isothiocyanate. **(b)** and (**c)** Modeled curves of dose response germination stimulant activities **(b)** and EC_50_ (half maximal effective concentration) **(c)**. Data presented ± SE. **(d**) and (**e)** Thermostability of PrKAI2d3 at 10 μM in the absence of ligand or in the presence of various ligands at 100 μM analyzed by nanoDSF. Changes in fluorescence (ratio F_350nm_/F_330nm_) with temperature (**d**) and first derivatives for the F_350nm_/F_330nm_ curve against the temperature gradient from which the apparent melting temperatures (Tm) were determined. for each sample (**e**). The experiment was carried out twice. **(f-h)** Mass spectrometry characterization of covalent PrKAI2d3-ligand complexes. Deconvoluted electrospray mass spectra of PrKAI2d3 (f) and PrKAI2d3^S98A^ (g) after addition of the 2-PEITC ligand, Peaks with an asterisk correspond to PrKAI2d3 covalently bound to a ligand (PrKAI2d3-ligand). The mass increments were measured for the PrKAI2d3-2-PEITC complex, 162.7 Da. Ligand-modified amino acids were identified by nanoLC-MSMS analyses after Glu-C proteolysis. (**h**) Fragmentation spectra of unmodified and different ligand-modified peptides. Labeled peaks correspond to the b and y fragments of the double-charged precursor ion displayed at the top. The PEITC ligand-modified serine residue is underlined. **(i)** Perception mechanism of PEITC.

As ITCs easily react with nucleophile functions^32^, we hypothesized that the PrKAI2d3 interaction with ITCs may trigger the formation of a covalent adduct. Indeed, a mass shift was detected correlating to the 2-PEITC covalently bound to the protein (Figure 4**f**). After the PrKAI2d3-2-PEITC complex digestion, the 2-PEITC attachment was localized on a peptide corresponding to the amino acids 87-113 of PrKAI2d3. Tandem mass spectrometry data revealed that the 2-PEITC attachment could be on His97 or on the catalytic Ser98 (Figure 4**h**). Incubation of 2-PEITC with PrKAI2d3^S98A^ allowed us to conclude that the major site for 2-PEITC attachment is on Ser98 (Figure 4**g**). These results hint at a perception mechanism for ITC in which the Ser98 hydroxyl group would react with isothiocyanate to generate a PrKAI2d3-attached carbamothioate (Figure 4**i**). However, no 2-PEITC attachment was detected on His249-containing peptides of the catalytic triad.

Overall, PrKAI2d3 acts as a receptor for both SL-like molecules and ITCs that are both germination stimulants of *P. ramosa*. These results demonstrate that further research into potential chemical interactors is achievable to design control methods.

### Search for small-molecule interactors with PrKAI2d3 proposes D-OH as a simple and efficient germination stimulant for *P. ramosa* with a bioactivity similar to that of ITCs

Recently, synthetic inhibitors of D14 and ShHTL7^33^, the *S. hermonthica* SL receptor, have been proposed, that include, although all structurally unrelated to SLs, soporidine, KK094, Triton X, and tolfenamic acid (TA) (Figure 5**a**). We evaluated the putative inhibitory effects of these compounds, along with those of a common serine protease inhibitor, phenylmethylsulfonyl fluoride (PMSF) on the germination of *P. ramosa* seeds treated with 10 nM of (±)-GR24 or 100 nM of 2-PEITC. Nonetheless, *P. ramosa* seeds were clearly hyposensitive to each of these compounds when compared to results obtained *in S. hermonthica*, with a maximum inhibition obtained in some instances at very high concentrations (Figure 5**b,c**; Supplementary Table 3). Indeed, all half maximum inhibitory concentrations (IC_50_) fell above the IC_50_ of abscisic acid (ABA, 100 nM (GR24) and 34 nM (2-PEITC)], a known inhibitor of *P. ramosa* germination^34,35^. These data indicate that the use of germination inhibitors intended for *Striga* is reasonably irrelevant and unsuitable for *P. ramosa* biological control.

**Figure 5.**
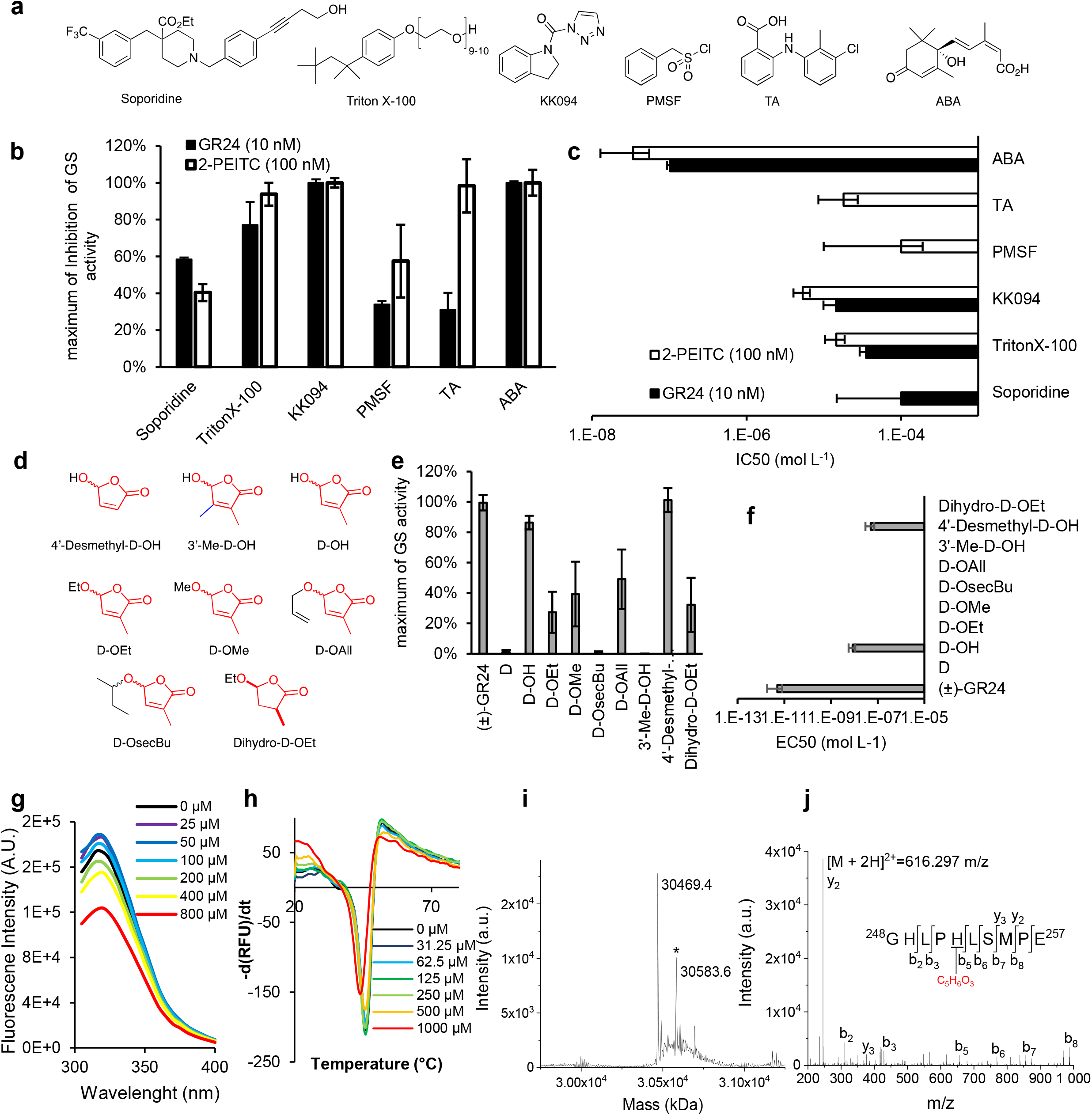
Identification of antagonists and agonists for the *P. ramosa* seed germination. **(a)** Chemical structures of inhibitors of D14 protein and absisic acid (ABA). (**b**) and (**c**) Comparison of the inhibitor effect on *P. ramosa* germination stimulation (GS) induced by GR24 (10 nM) or 2-PEITC (100 nM). Maximum of inhibition **(b)** and IC_50_ **(c)**. **(d)** Structure of D-OH rings and derivatives. **(e)** and **(f)** GS activity of D-OH and its derivatives on *P. ramosa* seeds. Maximum of GS activity (**e**) and EC_50_ (**f**). (**g**) D-OH binding of PrKAI2d3, based on intrinsic tryptophan fluorescence. Plots of fluorescence intensity *versus* probe concentrations. The change in intrinsic fluorescence was monitored and used to determine the apparent *K*_D_ values. The plots represent the mean of two replicates and the experiments were repeated at least three times. The analysis was done with GraphPad Prism 5.0 Software. **(h)** Biochemical analysis of the interaction between PrKAI2d3 at 10 μM and D-OH at various concentrations by DSF. Each line represents the average protein melt curve for two technical replicates and the experiment was carried out twice. **(i)** and (**j)** Mass spectrometry characterization of covalent PrKAI2d3-D-OH complexes. (**i**) Deconvoluted electrospray mass spectrum of PrKAI2d3 after addition of the D-OH ligand. Peaks with an asterisk correspond to PrKAI2d3 covalently bound to D-OH. The mass increments were measured for the PrKAI2d3-D-OH complex, 114.2 Da. Ligand-modified amino acids were identified by nanoLC-MSMS analyses after Glu-C proteolysis. (**j**) Fragmentation spectra of unmodified and different ligand-modified peptides. Labeled peaks correspond to the b and y fragments of the double-charged precursor ion displayed at the top. The D-OH ligand-modified histidine residue is underlined.

Therefore, we looked for specific PrKAI2d3 agonists with the potential for large-scale biocontrol. A serious issue in the search for simple and cheap SL analogs and mimics is their low stability due to the D-ring connection to a leaving group. As alternatives, molecules have been proposed that carry the D-ring only, because it is essential for the high bioactivity of SLs^36^. Specifically, the effect on the *P. ramosa* germination was evaluated of a set of butenolides containing the D-ring (D-OR) only (Figure 5**j**, Supplementary Figure 12). D-O*sec*Bu and 3’-Me-D-OH were completely inactive, whereas the *P. ramosa* sensitivity towards D-OAll, D-OMe, D-OEt, and Dihydro-DOEt was low with EC_50_ in the micromolar range. Inversely, D-OH and 4’-Desmethyl-D-OH were approximately 10-fold more active than 2-PEITC (Supplementary Table 2). Although D-OH had been found bioactive in rice (*Oryza sativa*) at high concentration^12^ (50 μM), in pea D-OH and 3’-Me-D-OH had not effect on the branching control by direct injection (100 μM) in the stem (Supplementary Figure 13).

To check whether this D-OH sensitivity was a specificity of the *P. ramosa* PrKAI2d3, we reexamined the germination-stimulating activity of D-OH on *S. hermonthica*. In contrast to previous results^37,38^, D-OH induced *S. hermonthica* germination (EC_50_ = 1.2 μM). Moreover, in *Arabidopsis* lines expressing a GFP fused to ShHTL7, the known *S. hermonthica* SL receptor^30,39^, was induced by (±)-DOH (EC_50_ = 3.1 μM) (Supplementary Figure 14, Supplementary Table 1). In summary, our study validates the perception of D-OH by SL receptors in root-parasitic plants.

DSF and intrinsic fluorescence analyses revealed that D-OH and PrKAI2d3 interacted at high concentrations (Figure 5**k-l**). Incubation of D-OH with PrKAI2d3 at pH 6.8 and the recorded MS spectra under denaturing conditions revealed a mass shift corresponding to the D-ring covalently bound to the protein (Figure 5**m**). The D-ring attachment could be localized to the His249 of the catalytic triad, similarly as for the SL analogs (Figure 5**n**). In addition to various SL analogs, and SL-like compounds, and ITCs, D-OH highlights the plasticity of the PrKAI2d3 receptor to interact with different structures with a significant biological activity as a consequence and emphasizes that all SL hydrolysis products are not inactive as germination stimulants of *P. ramosa*.

## Discussion

Obligate parasitic weeds require host-derived signals to germinate and wither their hosts long before they emerge from the soil, arguably making the early stages of the parasitic life cycle a much better target for control strategies than the later ones^40^. An important prerequisite for the design of such methods is an in-depth understanding of how parasites perceive germination stimulants.

Here, we demonstrated that PrKAI2d3 provides *P. ramosa* with hypersensitivity to SLs mainly due to its enzymatic activity. The enzymatic data with the GR24 analogs [(+)-GR24 and (−)-2’-*epi*-GR24] and GC probes suggest that PrKAI2d3 acts as a single turnover enzyme towards natural SLs, and cleave them to form a covalent complex between the D-ring and the catalytic histidine, as previously described for AtD14 and RMS3^13^. However, this model has recently been challenged^16^, because the *Arabidopsis* SL receptor mutated for the catalytic Asp residue, speculatively considered unable to cleave endogenous SLs, had been found to transduce the SL signal; hence, the SL cleavage has been concluded not be required for signaling. Indeed, It is possible that certain SL analogs, can be perceived independently of the enzymatic activity. We showed that the GR24 analogs with non-natural stereochemistry [(−)- GR24 and (+)-2’-*epi*-GR24] are not cleaved by PrKAI2d3. These uncleaved molecules stimulate 100-fold less efficiently the *P. ramosa* seed germination than the GR24 stereoisomers with a natural stereochemistry (Figure 3). Moreover, the PrKAI2d3 affinity (*K*_D_) towards the most bioactive SL analogs, (+)-GR24 and (−)-2’-*epi*-GR24, is in the micromolar range, which is several orders lower than the *P. ramosa* seed sensitivity recorded in the germination bioassays. The *K*_D_ values for (+)-GR24 and (−)-2’-*epi*-GR24, that reflect a simple binding to the receptor, do not allow explaining the hypersensitivity. Similar differences between *in vitro* and *in vivo* analyses were observed for the *S. hermonthica* SL receptor ShHTL7 characterization^30^. We propose that the SL hypersensitivity of this SL receptor is obtained by its enzymatic properties.

Interestingly, because the enzymatic activity is not completely abolished in the PrKAI2d3^S98A^ protein, a residually significant activity, also present in the DAD2^S96A^ protein^41^, might result from the nucleophilic addition of water instead of the attack of the Ser98 hydroxyl group. The partial complementation of the *htl-3* mutant with constructs that overproduce PrKAI2d3^S98A^ corroborates this assumption. A similar proposition could also explain why *Arabidopsis* overexpressing the AtD14 with a mutated Asp catalytic residue is able to transduce a branching inhibitor signal, especially the still untested natural SLs present in *Arabidopsis* (i.e. non-canonical SLs). Noteworthy, *p*-NPA could not efficiently highlight the enzymatic activity of PrKAI2d3, in comparison to the profluorescent SL mimics, pinpointing the weakness of using generic or inappropriate substrates as controls.

Moreover, PrKAI2d3 displays a high plasticity that allows binding of modified SLs, such as desmethyl-GR24 isomers or many SL mimics, but in contrast to RMS3 and AtD14, it perceives these compounds similarly as intact SLs. Correspondingly, desmethyl-YLG was bioactive in *Arabidopsis* via AtKAI2^42^. Ligand-mediated protein destabilization is not required to perceive SL analogs, but considerably improves the sensitivity to natural SLs. PrKAI2d3 acts as single-turnover enzyme for compounds with a D-ring without methyl group at the C4’ position, but a methyl group at the C-3’ position led to low interaction with PrKAI2d3. This result is in accordance with the low germination stimulant activity of 3’-methyl analogs and mimics for *P. ramosa* and corroborates studies conducted in *O. cumana*, *O. minor*, *P. aegyptiaca*, and *S. hermonthica*^28,43^. In other words, *P. ramosa* might be sensitive to D-ring modified SL and could perceive diverse SL-related compounds as putative SL degradation products^44^.

In addition, we established that ITCs, other known germination stimulants, interact with the PrKAI2d3 protein by forming a covalent complex. In *P. ramosa*, PrKAI2d3 can be considered as a germination stimulant receptor for different chemicals, including SLs, SL derivatives, and mimics, but also ITCs with completely different structures. Thus, *P. ramosa* has seemingly optimized its germination capacity by its ability to perceive various chemical mediators emitted by the host plants. The enzymatic activity-dependent ITC perception mechanism suggests that the active catalytic triad might have been conserved, not only to perceive SLs, but also other germination stimulants, not yet characterized for *P. ramosa*.

SLs are very unstable in the soil, especially under basic conditions that lead to ABC=CHOH and D-OH derivatives. To date, all SL analogs and mimics possess a D-ring connected to an ABC mimic by a hydrolysable function. The treatment with SL agonists developed until now induces a suicide germination of plant-parasitic seeds^45^. Nevertheless, these agonists are too unstable under field conditions to be effective, even when promising chemicals have been designed and applied for *S. hermonthica*^46^. In addition, cultures of P. *ramosa*-infested rapeseed mostly occur in slightly basic soils^47^, both favoring the formation of ITCs^4^ and complicating the use of SL mimics as a suicidal germination strategy, especially due to their instability. In sharp contrast, *P. ramosa* is highly sensitive to D-OH or DesMe-D-OH. As for D-H, no germination activity was detected, the hemiacetal function seems very important for the interaction with PrKAI2d3. Moreover, the residual bioactivity of D-OR (R ≠ H) might be explained by the hydrolysis of the R group during the bioassay, leading to D-OH. D-OH or DesMe-D-OH are especially interesting for translational research, because both compounds are small, are easily synthesized in comparison with synthetic SLs and SL mimics, and are not subjected to fast degradation. As such, D-OH and DesMe-D-OH seem to be promising chemicals for suicide germination in the field: their lower germination activity than that of SL mimics can be compensated by higher active concentration. Moreover, D-OH has the great advantage of being a natural product. The use of SL mimics, for which the part equivalent to the ABC moiety is in most cases xenobiotic, increases the risk of potential toxicity or pollution for the environment. In contrast to previous studies^37,38^, we found that D-OH is bioactive as a witchweed seed germination stimulant and that it could possibly be used for *S. hermonthica* biocontrol.

Here, we identified germination inhibitors and demonstrated the involvement of α/β-hydrolases for germination-stimulating reception, providing an alternative track to fight *P. ramosa*^48^. Novel more active inhibitors need to be designed for *P. ramosa* infection control without negative effects on the host plants and arbuscular mycorrhizal fungi. For this purpose, the identification of the SL receptor(s) in these fungi will undoubtedly be an important milestone. The use of specific inhibitors should be combined with other approaches for integrated management strategies^49^, such as chemical suicidal germination with compounds, such as D-OH, to decrease the parasitic-plant seed bank in soil.

## Methods

### Preparation of GR24 isomers, probes, and other ligands

For general experimental procedures, see the Supplementary Methods. (±)-2’-*epi*-GR24 and (±)-GR24 were prepared as described^50^ and (±)-4-desmethyl-GR24, (±)-2’-*epi*-4-desmethyl-GR24, and (±)-3’-Me-GR24 as described^27^. (+)-GR24, (−)-GR24, (+)-2’-*epi*-GR24, and (−)-2’-*epi*-GR24 were separated from (±)-2’-*epi*-GR24 and (±)-GR24 by chiral supercritical fluid chromatography as described^13,51^. (±)-GR24 was purified by semi-preparative HPLC by means of a Interchim puriFlash^®^ 4250 instrument, combined with a fraction collector with integrated ELSD, a PDA and a Phenomenex Luna C18, 250 × 21.2 mm, 5-μm column (H2O/CH3CN: 6/4) or Interchim Uptisphere Strategy SI, 250 × 21.2 mm, 5-μm column (Heptane/EtOAc: 1/1). 2-Phenethyl isothiocyanate (2-PEITC), benzylisothiocyanate (BEITC), DiFMU, *para*-nitrophenyl acetate (*p*-NPA), Yoshimulactone Green (YLG), abscisic acid (ABA), tolfenamic acid^52^ (TA), TritonX-100^53^ and phenylmethylsulfonyl fluoride (PMSF) are commercially available. KK094^54^ and soporidine^55^ were kindly provided by T. Asami and P. McCourt (University of Toronto), respectively. Probes (GC486, GC240, and GC242) were prepared as described^13^. For the preparations of D-OH and analogs, see Supplementary Methods.

### Expression and purifications

AtD14 and AtKAI2 were purified and expressed with cleavable GST tags as described^13^. For PrKAI2d3 expression, the coding sequences from *Phelipanche ramosa* were amplified by PCR by means of a seed-derived cDNA template and specific primers (Supplementary Table 4) containing a protease cleavage site for tag removal, and subsequently cloned into the pGEXT-4T-3 expression vector. The PrKAI2d3 and PrKAI2d3^S98A^ proteins were purified and expressed as above.

### Site-directed mutagenesis

Site-directed mutagenesis experiments were done with the QuickChange II XL Site Directed Mutagenesis kit (Stratagene) on pGEX-4T-3-PrKAI2d3 (Supplementary Table 4). Mutagenesis was verified by systematic DNA sequencing.

### Plant material and growth conditions

Pea (*Pisum sativum*) branching mutant plants were derived from various cultivars after ethyl methanesulfonate (EMS) mutagenesis and had been described previously^56^. The *rms1-10* (M3T-884) mutant was obtained from the dwarf cv Térèse. Plants were grown in a greenhouse under long-day as described^57^.

All *Arabidopsis thaliana* (L.) Heynh. mutant plants (Columbia-0 [Col-0] accession background) have been described previously: *htl-3* (ref. ^58,59^), *Atd14-1/htl-3* (ref. ^58,59^), and *htl-3* ShHTL7 (kind gift of P. McCourt). For overexpression of the green fluorescent protein (GFP) fusions of PrKAI2d3 and PrKAI2d3^S98A^, transgenic *Arabidopsis* seeds were generated by the *Agrobacterium* floral dip method^60^ with the *htl-3* mutant as the background accession. Transgenic seeds were selected based on the antibiotic resistance and GFP fluorescence.

Two batches of parasitic plant seeds were used. A population of seeds of *Phelipanche ramosa* (L.) Pomel associated to the genetic group 1 (*P. ramosa* 1) was collected from Saint Martin-de-Fraigneau (France) on broomrape-parasitizing winter rapeseed (*Brassica napus* L.) in 2014 and 2015 (ref. ^61^). Seeds of *Striga hermonthica* (Delile) Benth. (Sudan, 2007) were provided by Lukas Spichal (Olomuc, Czech Republic). Seeds were surface sterilized and conditioned as described^29^ (darkness; 21 °C and 30 °C for *P. ramosa* and *S. hermonthica*, respectively).

### Pea shoot-branching assay

The compounds to be tested were applied by vascular supply. The control was the treatment with 0.1% dimethylsulfoxide (DMSO) only. Twelve plants were sown per treatment in trays and generally 10 days after sowing the axillary bud at node 3 was treated. Compounds in DMSO solution were diluted in water to the indicated concentrations for a treatment with 0.1% (v/v) DMSO. The branches at nodes 1 and 2 were removed to encourage the outgrowth of axillary buds at the nodes above. Nodes were numbered acropetally from the first scale leaf as node 1 and cotyledonary node as node 0. Bud growth at nodes 3 and 4 was measured with digital callipers 8 to 10 days after treatment. Plants with damaged main shoot apex or with a dead white treated-bud were discarded from the analysis. The SL-deficient *rms1-10* pea mutant was used for all experiments.

### Cloning and generation of transgenic lines

For all GFP fusion constructs, cloning was done by Gateway recombination (Thermo Fisher Scientific). The open reading frame (ORF) of PrKAI2d3 was amplified from *P. ramosa* cDNA with iProof™ High-Fidelity DNA Polymerase (Bio-Rad) and the Gateway^®^-specific primers PrKAI2d3_B2_FW and PrKAI2d3_B3_Rev_STOP. The PCR product flanked by the attB sites was cloned in pDONR P2R-P3 with the BP Clonase II enzyme mix (Invitrogen). The resulting entry vector was used to clone the genes into the destination vector pK7m34GW, under the control of the 35S promoter, and N-terminally fused with GFP with the LR Clonase II Plus enzyme mix (Invitrogen). For the construction of the catalytic site mutant, pDONR P2R-P3-PrKAI2d3 was mutated with the QuikChange II site directed mutagenesis kit (Agilent). The generated clones were checked by sequencing. All primers used for cloning are listed (Supplementary Table 4).

### Western Blot

Total protein was extracted from 5-day-old seedlings, exposed to white light for 3 h, transferred to darkness for 21 h, and exposed to continuous red light for 4 days. Protein concentrations were determined by the Bradford assay (Bio-Rad). Of the protein extracts, separated by sodium dodecyl sulfate-polyacrylamide gel electrophoresis (SDS-PAGE) and transferred onto polyvinylidene fluoride (PVDF) membranes, 30 μg was detected with horseradish peroxidase (HRP)-conjugated antibodies against GFP (Anti-GFP-HRP, 1:10000, Miltenyi Biotec) or anti-tubulin (mouse monoclonal, 1/10000, Sigma-Aldrich) and HRP-conjugated anti-mouse antibodies (rabbit polyclonal, 1/10000, Abcam). The blots were visualized with the Western Lightning Plus Enhanced Chemiluminescence kit (PerkinElmer) and the X-Doc System (Bio-Rad). The Precision Plus Protein™ Dual Color Standards (Bio-Rad) was used as protein size marker.

### *Arabidopsis* hypocotyl elongation assays

*Arabidopsis* seeds were surface sterilized by consecutive treatments of 5 min 70% (v/v) ethanol with 0.05% (w/v) SDS and 5 min 95% (v/v) ethanol and sown on half-strength Murashige and Skoog (½MS) media (Duchefa Biochemie) containing 1% (w/v) agar, supplemented with 1 μM (±)-GR24 [0.01% (v/v) DMSO] or with 0.01% (v/v) DMSO only (control). Seeds were stratified at 4 °C for 2 days in the dark, then exposed to white light for 3 h, transferred to darkness for 21 h, and exposed to continuous red light for 4 days at 21°C. Plates were photographed and hypocotyl lengths were quantified using ImageJ (http://imagej.nih.gov/ij/).

### *Arabidopsis* germination assays

*Arabidopsis* seeds were after-ripened for at least 6 weeks before use. Surface-sterilized seeds were incubated in the incubation solution (1 mM HEPES buffer; pH 7.5) at a ratio of 10 mg of seeds per mL^35^. Fifty μL of seeds (~ 20-25 seeds) were distributed on 96-well plates and 10 μL of germination stimulants [(10 μM (+)-GR24, 10 μM (−)-GR24, and 0.1% (v/v) DMSO (control) or 10-fold concentrated D-OH] were added. The final volume was adjusted to 100 μL with the incubation solution. Plates were incubated either for 5 days at 25 °C in the dark^62^ or for 7 days at 32 °C-34 °C under constant light illumination at a quantum irradiance of 10 μmol m^−2^ s^−1^ (ref. ^20,26^). A seed was considered germinated when the radicle protruded from the seed coat.

### Germination stimulation activity assay on root-parasitic plant seeds

Germination stimulant activity of chemicals on seeds of parasitic plants were determined as described previously^29^. Chemicals were suspended in DMSO at 10 mmol L^−1^, diluted with water at 1 mmol L^−1^ (water/DMSO; v/v; 9/1), and then of 1×10^−5^ mol L^−1^ to 1×10^−12^ mol L^−1^ with water/DMSO (v/v; 9/1). For each compound, a concentration range from 10^−13^ to 10^−6^ mol L^−1^ (water/DMSO; 99/1) was applied to the conditioned parasitic seeds. As negative control, 1% (v/v) DMSO was used (seed germination < 1%) and as positive control (±)-GR24 at a concentration of 1 μmol L^−1^, inducing 72-87% and 50–65% of seed germination for *P. ramosa* 1 and *S. hermonthica*, respectively. To avoid variations related to sterilization events, the germination percentages are reported as a ratio relative to the positive control [1 μmol L^−1^ (±)-GR24) included in each germination assay. Each dilution and germination assay were repeated at least three times. For each compound tested, dose response curves (germination stimulation activity = f(c); germination stimulant activity relative to 1 μmol L^−1^ (±)-GR24; c, concentration (mol L^−1^); half maximal effective concentration (EC_50_); and maximum germination stimulant activity) were determined with a Four Parameter Logistic Curve computed with SigmaPlot^®^ 10.0.

### Enzymatic degradation of GR24 isomers by purified proteins

The ligand (10 μM) was incubated without and with purified AtD14/AtKAI2/PrKAI2d3/PrKAI2d3^S98^A (5 μM) for 150 min at 25°C in 0.1 mL phosphate buffered saline (PBS; 100 mM Phosphate, pH 6.8, 150 mM NaCl) in the presence of (±)-1-indanol (100 M) as internal standard. The solutions were acidified to pH = 1 by addition of 2 μL trifluoroacetic acid (TFA) to quench the reaction and centrifugated (12 min, 12,000 tr/min). Thereafter, the samples were subjected to reverse-phase-ultra-performance liquid chromatography (RP-UPLC)-MS analyses by means of UPLC system equipped with a Photo Diode Array (PDA) and a Triple Quadrupole Detector (TQD) mass spectrometer (Acquity UPLC-TQD, Waters). RP-UPLC (HSS C_18_ column, 1.8 μm, 2.1 mm × 50 mm) with 0.1% (v/v) formic acid in CH_3_CN and 0.1% (v/v) formic acid in water (aq. FA, 0.1%, v/v, pH 2.8) as eluents [10% CH_3_CN, followed by linear gradient from 10% to 100% of CH_3_CN (4 min)] at a flow rate of 0.6 mL min^−1^. The detection was done by PDA and with the TQD mass spectrometer operated in Electrospray ionization-positive mode at 3.2 kV capillary voltage. To maximize the signal, the cone voltage and collision energy were optimized to 20 V and 12 eV, respectively. The collision gas was argon at a pressure maintained near 4.5 10^−3^ mBar.

### Enzymatic assays with profluorescent probes and *p*-nitrophenyl acetate

The assays were done as described^13^ with a TriStar LB 941 Multimode Microplate Reader (Berthold Technologies).

### Protein melting temperatures

For the Differential Scanning Fluorimetry (DSF) experiments a CFX96 Touch^™^ Real-Time PCR Detection System (Bio-Rad) was used with excitation and emission wavelengths of 490 and 575 nm, respectively and Sypro Orange (λex/λem: 470/570 nm; Life Technologies) as the reporter dye. Samples were heat-denatured with a linear 25 °C to 95 °C gradient at a rate of 1.3 °C per min after incubation at 25 °C for 30 min in the dark. The denaturation curve was obtained by means of the CFX manager™ software. Final reaction mixtures were prepared in triplicate in 96-well white microplates. Each reaction was carried out in 20-μL sample in PBS (100 mM phosphate, pH 6.8, 150 mM NaCl) containing 6 μg of protein (so that the final reactions contained 10 μM protein), 0 to X ligand concentrations in μM, 4% (v/v) DMSO, and 0.008 μL Sypro Orange. Plates were incubated in the dark for 30 min before analysis. In the control reaction, DMSO was added instead of the chemical solution. The experiments were repeated three times.

### nanoDSF

Proteins were diluted in PBS (100 mM Phosphate, pH 6.8, 150 mM NaCl) to a concentration of ~10 μM. Ligands were tested at the concentration of 200 μM. The intrinsic fluorescence signal was measured as a function of increasing temperature with a Prometheus NT.48 fluorimeter (NanoTemper Technologies), with 55% excitation light intensity and 1 °C/min temperature ramp. Analyses were done on capillaries filled with 10 μL of the respective samples. Intrinsic fluorescence signals expressed by the 350 nm/330 nm emission ratio that increases as the proteins unfold, were plotted as a function of temperature (Figures **2d**, **2f,** and **4d**). The plots show one of the three independent data collections done for each protein.

### Intrinsic tryptophan fluorescence assays and determination of the dissociation constant *K*_D_

The experiments were done as described^13^ by means of the Spark^®^ Multimode Microplate Reader (Tecan).

### Direct electrospray ionization (ESI)-MS under denaturing conditions

Mass spectrometry measurements were carried out with an electrospray quadrupole-time of flight (Q-TOF) mass spectrometer (Waters) equipped with the Nanomate device (Advion). The HD_A_384 chip (5 μm i.d. nozzle chip, flow rate range 100-500 nL/min) was calibrated before use. For the ESI-MS measurements, the Q-TOF instrument was operated in radio frequency quadrupole mode with the TOF data collected between *m/z* 400-2990. The collision energy was set to 10 eV and argon was used as collision gas. Mass spectra were acquired after denaturation of PrKAI2d3 ± ligand in 50% (v/v) acetonitrile and 1% (v/v) formic acid. The Mass Lynx 4.1 (Waters) and Peakview 2.2 (AB Sciex) softwares were used for data acquisition and processing, respectively. Multiply-charged ions were deconvoluted by the MaxEnt algorithm (AB Sciex). The protein average masses were annotated in the spectra and the estimated mass accuracy was ± 2 Da. For the external calibration, NaI clusters (2 μg/μL, isopropanol/H2O 50/50, Waters) were used in the acquisition *m/z* mass range.

### Localization of the fixation site of ligands on PrKAI2d3

PrKAI2d3-ligand mixtures were incubated for 10 min prior overnight Glu-C proteolysis. Glu-C--generated peptide mixtures were analyzed by nanoLC-MS/MS with the Triple-TOF 4600 mass spectrometer (AB Sciex) coupled to the nanoRSLC UPLC system (Thermo Fisher Scientific) equipped with a trap column (Acclaim PepMap 100 C_18_, 75 μm i.d. × 2 cm, 3 μm) and an analytical column (Acclaim PepMap RSLC C_18_, 75 μm i.d. × 25 cm, 2 μm, 100 Å). Peptides were loaded at 5 μL/min with 0.05% (v/v) TFA in 5% (v/v) acetonitrile and separated at a flow rate of 300 nL/min with a 5% to 35% solvent B [0.1% (v/v) formic acid in 100% acetonitrile] gradient in 40 min with solvent A [0.1% (v/v) formic acid in water]. NanoLC-MS/MS experiments were conducted in a Data-Dependent acquisition method by selecting the 20 most intense precursors for collision-induced dissociation fragmentation with the Q1 quadrupole set at a low resolution for better sensitivity. Raw data were processed with the MS Data Converter tool (AB Sciex) for generation of .mgf data files and proteins were identified with the MASCOT search engine (Matrix Science) against the PrKAI2 sequence with oxidation of methionines and ligand-histidine adduct as variable modifications. Peptide and fragment tolerance were set at 20 ppm and 0.05 Da, respectively. Only peptides were considered with a MASCOT ion score above the identity threshold (25) calculated at 1% false discovery rate.

### Phylogenetic analysis

Phylogenetic analyses were done on 32 D14 and KAI2 sequences, containing five sequences of *Phelipanche ramosa* obtained in this study and previously described sequences from *Phelipanche aegyptiaca, Striga hermonthica, Pisum sativum, Arabidopsis thaliana*, and *Oryza sativa*^13,17,19,20,63,64^. The D14 and KAI2 proteins and the nucleotide data (Supplementary Data files S1 and S2, respectively) were aligned with AFFT^65^ and the G-INS-i iterative refinement alignment method^66^. Sequence alignments (Supplementary Data files S3 and S4) were manually trimmed to remove gaps at either end, producing final protein and nucleotide data sets of 262-264 amino acids and 786-792 nucleotides, respectively (Supplementary Data files S5 and S6). The α/β hydrolase RbsQ from *Bacillus subtilis* was used as outgroup, because of its high similarity to KAI2 and D14, and a conserved catalytic triad^17^. Maximum likelihood (ML) analyses were conducted with RAxML^67^ on 1,000 bootstraps replicates for statistical support of our inferences. The best ML tree for the amino acid sequences was inferred with the PROT-GAMMA model and the WAG substitution matrix. For the nucleotides sequences, the best ML tree was searched with the GTR model and a gamma rate of heterogeneity among sites. The percentage of trees in which the associated taxa clustered together was provided next to the branches. The resulting consensus amino acid and nucleotide trees were drawn to scale with MEGA7 (ref. ^68^), with the branch lengths representing substitutions per sites.

### Modeling

Protein sequences of the *P. ramosa* KAI2 proteins were modeled by means of the SWISSMODEL server (http://swissmodel.expasy.org/)^69^. Models were generated with the chain A of the Apo form of the *Arabidopsis* KAI2 structure (PDB: 4JYP) as a template^70^. Figures of protein structure were generated with PYMOL. Cavities within homology models were visualized by the surface mode on the setting “cavities and pockets culled” within PYMOL. Pocket sizes were calculated with the CASTp 3.0 server^71^ and a probe radius of 1.4 Å. The reported pocket sizes were the Connolly’ solvent excluded surface volumes of the catalytic pocket.

### Expression data

For the complemented *Arabidopsis* lines, seeds were imbibed for 24 h in a growth solution (1 mM HEPES buffer, pH 7.5, 0.1 mM Preservative Plant Mixture in sterile water) and treated for 6 h with 10 μM (±)-GR24 or mock solution. *P. ramosa* seeds were imbibed in the growth solution for 7 days and then treated for 1 h with 1 nM (±)-GR24 or mock solution. Seeds were harvested, snap-frozen in liquid nitrogen, and ground with pestle and mortar to a fine powder. RNA was extracted and purified with the RNeasy Plant mini kit (Qiagen). Genomic DNA was removed by DNase treatment and the samples were purified by ammonium acetate (5 M final concentration) precipitation. The iScript cDNA synthesis kit (Bio-Rad) was used to reverse transcribe RNA. SYBR Green detection was used during qRT-PCR run on a Light Cycler 480 (Roche). Reactions were done in triplicate in a 384-multiwell plate, in a total volume of 5 μL and a cDNA fraction of 10%. Cycle threshold values were obtained and analyzed with the 2-ΔΔCT method^72^. The values from three biological and three technical repeats were normalized against those of the seed-specific housekeeping gene At4g12590 for *Arabidopsis*^73^. For *P. ramosa, PrCACS* was used to normalize the expressions. The normalized values were analyzed according to the published model^74^ with the mixed model procedure (SAS Enterprise).

### Statistical analyses

As deviations from normality had been observed for the axillary bud length, hypocotyl length, and germination after SL treatments, the Kruskal-Wallis test was used to assess the significance of one treatment with one compound in comparison to a treatment with another by means of the R Commander software (version 1.7–3) (ref.^75^).

## Aknowledgements

We thank J.-P. Pillot for the pea plant bioassays, Bruno Baron (Institut Pasteur, France) for access to and help with the nanoDSF experiments, Thomas Larribeau for technical assistance, and Catherine Rameau, Sandrine Bonhomme and Martine De Cock for their comments on the manuscript. This work was supported by the Institut Jean-Pierre Bourgin’s Plant Observatory technological platforms, a “Infrastructures en Biologie Santé et Agronomie” grant to SICAPS platform of the Institute for Integrative Biology of the Cell, and CHARM3AT Labex program (ANR-11-LABX-39). A.d.S.G. is the recipient of an AgreenSkills award from the European Union in the framework of the Marie-Curie FP7 COFUND People Programme and fellowship from Saclay Plant Sciences (ANR-17-EUR-0007). A.J. is indebted to the Research Foundation-Flanders for a Structural Basic Research fellowship (Project 1S15817N) and for a travel grant in the framework of a Tournesol fellowship (Project VS04418N).

## Author contributions

A.d.S.G., J.-B.P., and F.-D.B. designed research; G.C. designed and synthesized the profluorescent probes; V.Se. and F.-D.B. synthesized the chemicals; A.d.S.G., A.J., and E.B. produced and purified the proteins; A.d.S.G. and A.J. characterized the proteins and did the kinetic experiments; A.J., G.B., J.-B.P., and F.-D.B. performed the plant experiments; D.C. performed the mass experiments; A.d.S.G., V.St., and F.-D.B. performed the HPLC analyses and separations; A.d.S.G., A.J., G.B., J.-B.P., L.B, D.C., K.G., P.S., S.W., S.G., P.D., and F.-D.B. analyzed the data; A.d.S.G., A.J., G.B., J.-B.P., and F.-D.B. wrote the paper. All authors critically revised the manuscript.

## Additional information

**Supplementary information** is available for this paper at www…………….

**Materials and Correspondence and requests for materials** should be addressed to F.-D.B.

## Supplementary information

Supplementary Figure 1. **Chemical structures. (a)** Natural strigolactones. **(b)** Profluorescent probes. GC series and Yoshimulactone (YLG).

Supplementary Figure 2. **Phylogenetic analysis of KAI2 and D14 nucleotide sequences.** The phylogenetic tree was constructed with the maximum likelihood method and 1,000 bootstraps replicates by means of RAxML. Scale bar = 0.1 substitutions per site.

Supplementary Figure 3. **Sequence alignment of *Phelipanche ramosa* (*Pr*) and *P. aegyptiaca* (*Pa*) KAI2 protein with D14 and KAI2 proteins from *Arabidopsis* (At), rice (Os), pea (RMS), and the bacterial RbsQ.** The three amino acid residues corresponding to the catalytic triad are marked with asterisks. Amino residues highlighted in the Figure 1**b** are indicated with a blue arrowhead. Amino acid numbers are indicated for AtD14. Note that the rice OsD14 protein has a non-conserved 50-amino-acid N-terminal extension omitted in the alignment.

Supplementary Figure 4. **Dose response germination stimulant activities of (+)-GR24 and (−)-GR24 with *Arabidopsis* lines.** Dose response germination stimulation activities and modeled curves of (+)-GR24 **(a)** and (−)-GR24 **(b)** on seeds of *Arabidopsis* Col-0 and *htl-3*/p35S::PrKAI2d3. **(c)** Half maximal effective concentration (EC_50_). **(d)** Maximum germination stimulation activities. Data are indicated ± SE. **(e)** Relative accumulation of *PrKAI2d3* transcripts in *Arabidopsis* Col-0, *htl-3*, and p35S::GFP-PrKAI2d3 #4 (*htl-3*) seeds imbibed for 24 h. **(f)** Accumulation of PrKAI2d3 proteins in *Arabidopsis* p35S::GFP-PrKAI2d3 #4 (*htl-3*) and *htl-3* 5d-old seedlings.

Supplementary Figure 5. **Germination assay on *P. ramosa* parasitic-plant seeds of GR24 isomers [(+)-GR24, (−)-GR24, (+)-2’-*epi*-GR24, and (−)-2’-*epi*-GR24] and methyl variation on the GR24 D-ring [(±)-Desmethyl-GR24, (±)-4’-Desmethyl-2’-epi-GR24, and (±)-3’-Me-GR24]**. **(a**) and **(c)** Dose response germination stimulation activities and modeled curves.**(b)** and (**d)** Maximum germination stimulation activity relative to (±)-GR24 (1 μM). Data are indicated ± SE.

Supplementary Figure 6. **Germination assay on *P. ramosa* parasitic-plant seeds of various profluorescent ligands and biochemical analysis of the interaction between PrKAI2d3 and various ligands by DSF. (a)** Dose response germination stimulation activities and modeled curves. **(b)** Maximum germination stimulant activity relative to (±)-GR24 (1 μM). **(c)** Half maximal effective concentration (EC_50_) (mol L^−1^). Data are indicated ± SE. **(d-i)** Melting temperature curves of PrKAI2d3 with (±)-GR24 **(d)**, (±)-GC486 **(e)**, (±)-GC240 **(f)**, DiFMU **(g)**, (+)-GC242 **(h),** and (−)-GC242 **(i)** at varying concentrations assessed by DSF. Each line represents the average protein melt curve for three technical replicates and the experiment was carried out twice.

Supplementary Figure 7. **Biochemical analysis of the interaction between PrKAI2d3 (a-d, i-k) or PrKAI2d3^S98A^ (e-h, l-n) and various ligands by DSF.** Melting temperature curves of PrKAI2d3 or PrKAI2d3^S98A^ at 10 μM with (+)-GR24 **(a, e)**, (−)-GR24 **(b, f)**, (+)-2’-*epi*-GR24 **(c, g)**, (−)-2’-*epi*-GR24 **(d, h)**, (±)-GR24 **(i, l)**, (±)-4’-desmethyl-2’-*epi*-GR24 **(j, m)**, or (±)-3’-Me-GR24 **(k, n)** at varying concentrations assessed by DSF. Each line represents the average protein melt curve for three technical replicates and the experiment was carried out twice.

Supplementary Figure 8. **Intrinsic tryptophan fluorescence of PrKAI2d3 in the presence of SL analogs.** Changes in intrinsic fluorescence emission spectra of PrKAI2d3 in the presence of various concentrations of (+)-GR24 **(a)**, (−)-GR24 **(b)**, (+)-2’-*epi*-GR24 **(c)**, (−)- *2’-epi-GR24* **(d)**, (±)-GR24 **(e)**, (±)-4’-desmethyl-2’-*epi*-GR24 **(f)**, or (±)-3’-Me-GR24 **(g)**. Proteins (10 μM) were incubated with increasing amounts of ligand (0–800 μM from top to bottom). The observed relative changes in intrinsic fluorescence were plotted as a function of the SL analog concentration, transformed to a saturation degree, and used to determine the apparent *K*D values relevant to Figure 2**h,i**. The plots represent the mean of two replicates and the experiments were repeated at least three times. The analysis was done with GraphPad Prism 8.0 Software.

Supplementary Figure 9. **Intrinsic tryptophan fluorescence of PrKAI2d3 in the presence of profluorescent SL probes.** Changes in intrinsic fluorescence emission spectra of PrKAI2d3 in the presence of various concentrations of (±)-GC486 (**a**), (±)-GC240 (**b**), DiFMU (**c**), YLG (**d**), (±)-GC242 (**i**), (+)-GC242 (**j**), or (−)-GC242 (**k**). Proteins (10 μM) were incubated with increasing amounts of ligand (0–800 μM from top to bottom). The observed relative changes in intrinsic fluorescence were plotted as a function of the SL analog concentration and transformed to saturation. Plots of fluorescence intensity *versus* (±)-GC486 (**e**), (±)-GC240 (**f**), DiFMU (**g**), and YLG concentrations used to determine the apparent *K*D values. The plots represent the mean of two replicates and the experiments were repeated at least three times. The analysis was done with GraphPad Prism 8.0 Software.

Supplementary Figure 10. **PrKAI2d3 hydrolysis activity.** Progress curves during the 4-nitrophenyl acetate (*p*-NPA) (1 mM) hydrolysis by PrKAI2d3, PrKAI2d3^S98A^, AtKAI2, and AtD14 (4 μM). The *p*-NPA release was monitored (A_405_) at 25 °C.

Supplementary Figure 11. **Biochemical analysis of the interaction between PrKAI2d3 and ITCs by DSF.** Melting temperature curves of PrKAI2d3 at 10 μM with (±)-GR24 **(a)**, 2-PEITC **(b)**, or BITC **(c)** at varying concentrations assessed by DSF. Each line represents the average protein melt curve for three technical replicates and the experiment was carried out twice.

Supplementary Figure 12. **Germination inhibition by various chemicals and stimulation by D-analogs.** Dose response germination stimulation (GS) activities with 10 nM GR24 **(a)** and 100 nM 2-PEITC (**b)** and modeled curves **(c**) and (**d)**. Maximum of germination stimulation activity relative to (±)-GR24 (1 μM). Data are indicated ± SE.

Supplementary Figure 13. **Bud outgrowth inhibition activity assay for D derivatives after direct stem infusion.** Data are means ± SE (*n* = 12), 8 days after treatment of the pea plants *rms1-10* and Térèse as control. \**P* < 0.5, \*\*\**P* < 0.001, Kruskal-Wallis rank sum test, compared to control values (CTL0).

Supplementary Figure 14. **Perception of the germination stimulant D-OH by ShHTL7.** Dose response germination stimulation activities and modeled curves of (±)-GR24 and D-OH on seeds of *S. hermonthica* **(a),** *Arabidopsis* Col-0 (**c)**, and *Arabidopsis htl-3*, ShHTL7 (34 °C, continuous light for *Arabidopsis* (**d)**. **(b)** Seed germination of *Arabidopsis* Col-0, *htl-3, htl-3*/ShHTL7, and *max2* with GR24 and D-OH (10 μM). **(e)** Half maximal effective concentration (EC_50_). **(f)** Maximum germination stimulation activities. Data are indicated ± SE.

Supplementary Table 1. EC_50_ and maximum germination percentage of (+)-GR24, and (−)- GR24 and of (±)-GR24 and D-OH in *Arabidopsis* lines.

Supplementary Table 2. EC_50_ and maximum of germination stimulant activity of compounds for *P. ramosa*.

Supplementary Table 3. IC_50_ and maximum of inhibition of putative inhibitors of PrKAI2d3 for *P. ramosa*.

Supplementary Table 4. Oligonucleotides used.

