## Supplementary Figures and Tables for "A *Phelipanche ramosa* KAI2 Protein Perceives enzymatically Strigolactones and Isothiocyanates"

Alexandre de Saint Germain,[a] Anse Jacobs,[b-e] Guillaume Brun,[f,g] Jean-Bernard Pouvreau,[f] Lukas Braem,[b-e] David Cornu,[h] Guillaume Clavé,[i] Emmanuelle Baudu,[a] Vincent Steinmetz,[i] Vincent Servajean,[i] Susann Wicke,[g] Kris Gevaert,[d,e] Philippe Simier,[f] Sofie Goormachtig,[b,c] Philippe Delavault[f] and François-Didier Boyer[i]*

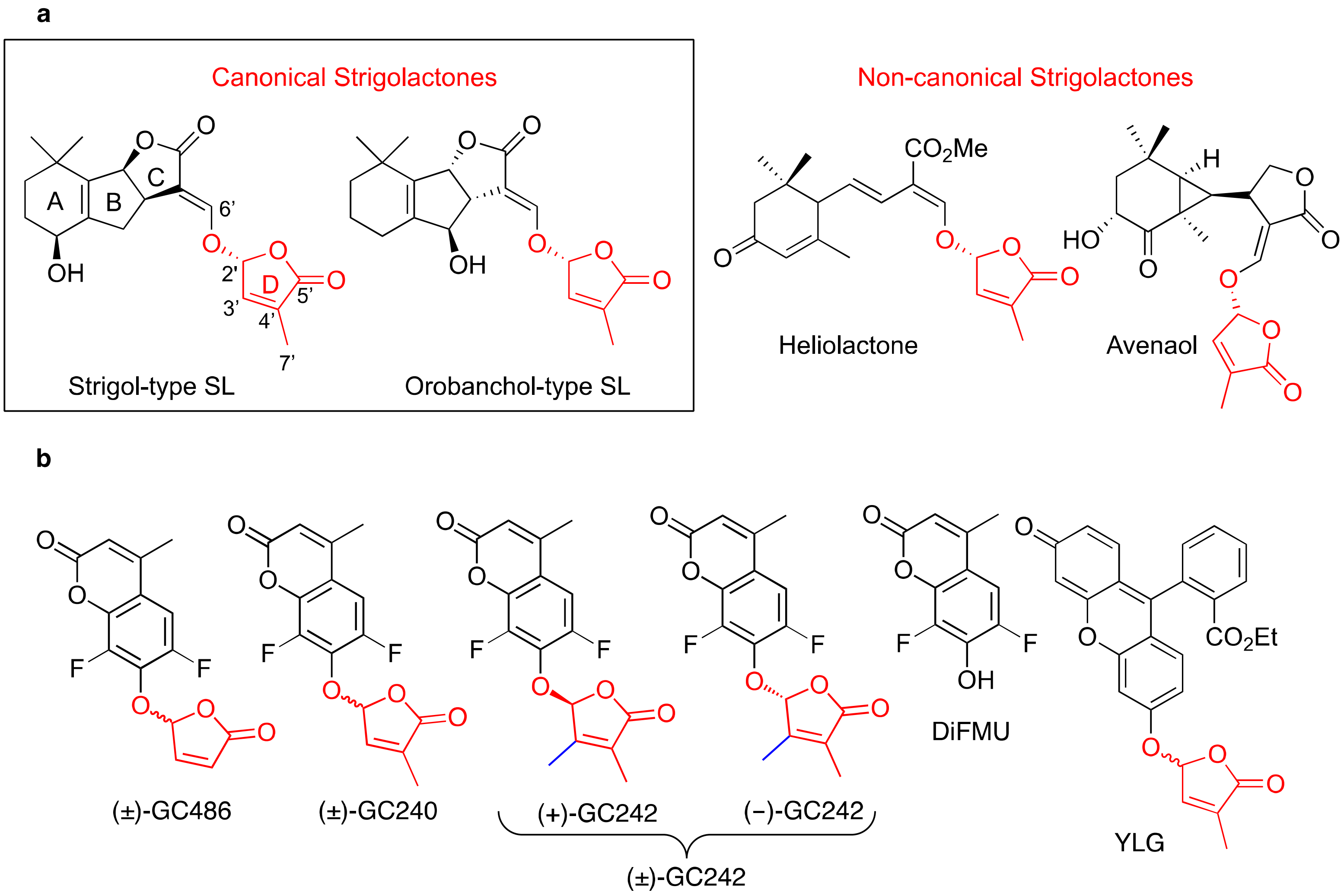

**Supplementary Figure 1.**

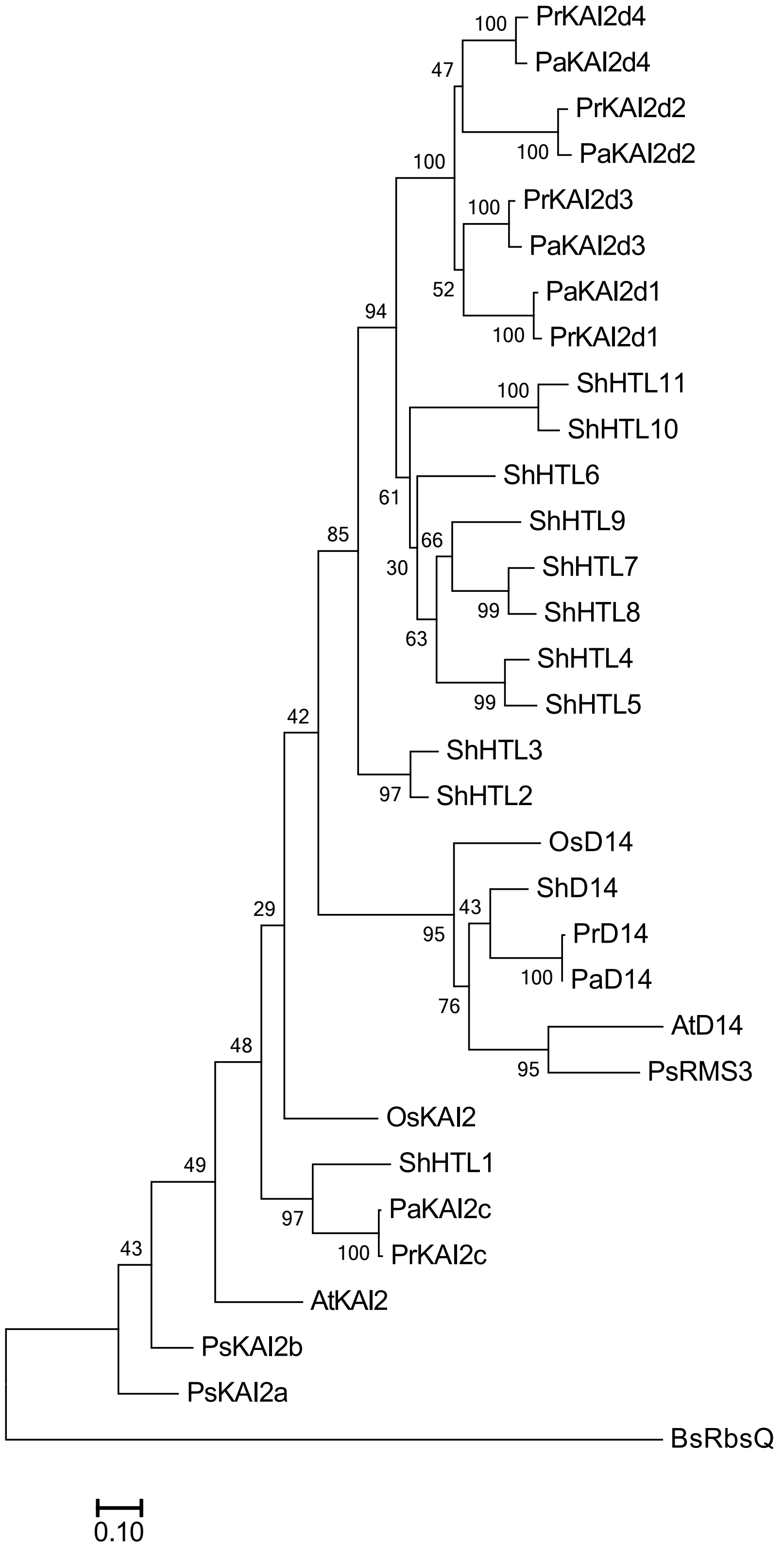

**Supplementary Figure 2.**

Data file S1. D14 and KAI2 amino acid sequences
Data file S2. D14 and KAI2 nucleotide sequences
Data file S3. MAFFT alignement of D14 and KAI2 amino acid sequences
Data file S4. MAFFT alignement of D14 and KAI2 nucleotide sequences
Data file S5. Trimmed MAFFT alignement of D14 and KAI2 amino acid sequences
Data file S6. Trimmed MAFFT alignement of D14 and KAI2 nucleotide sequences

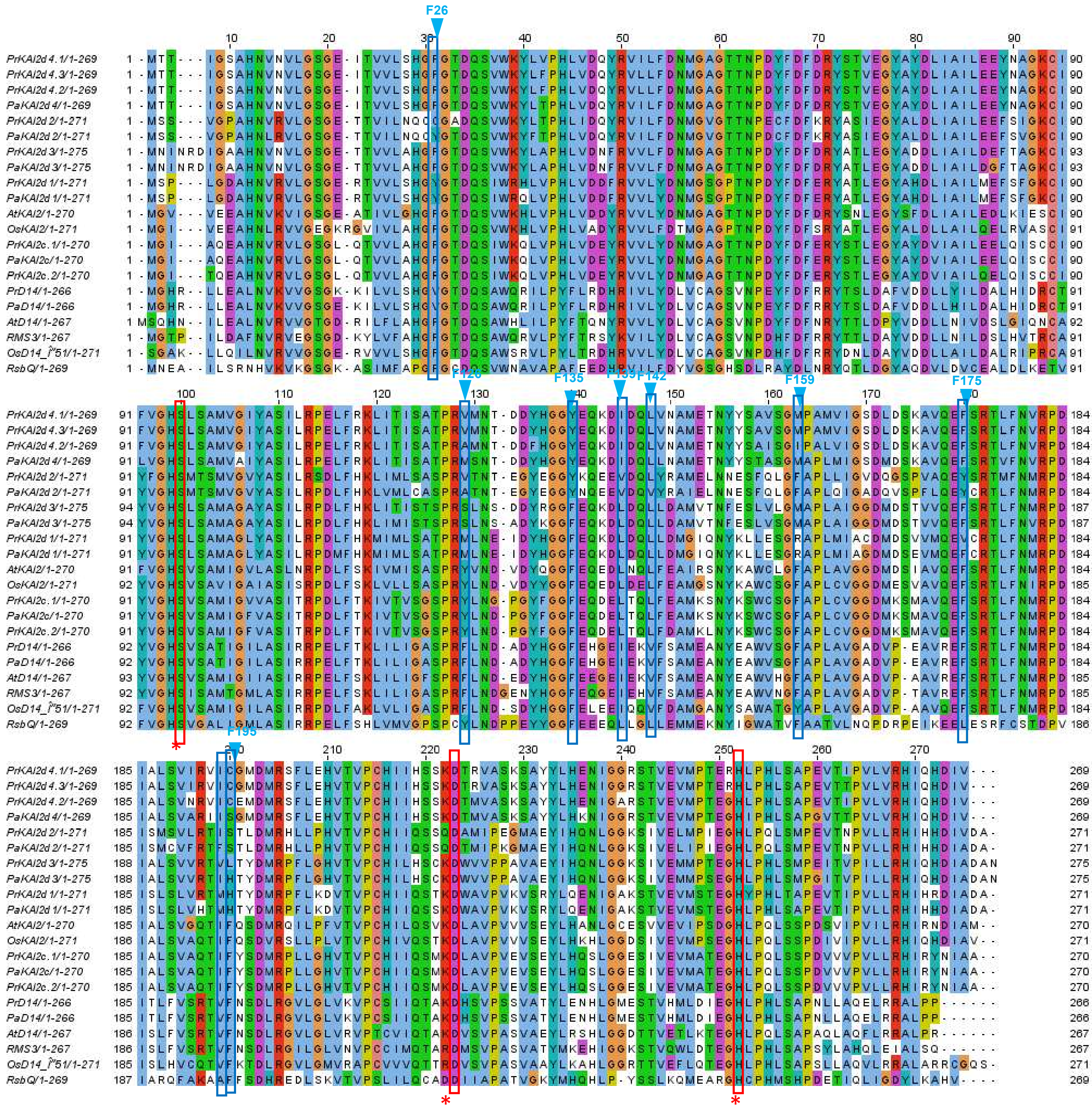

**Supplementary Figure 3.**

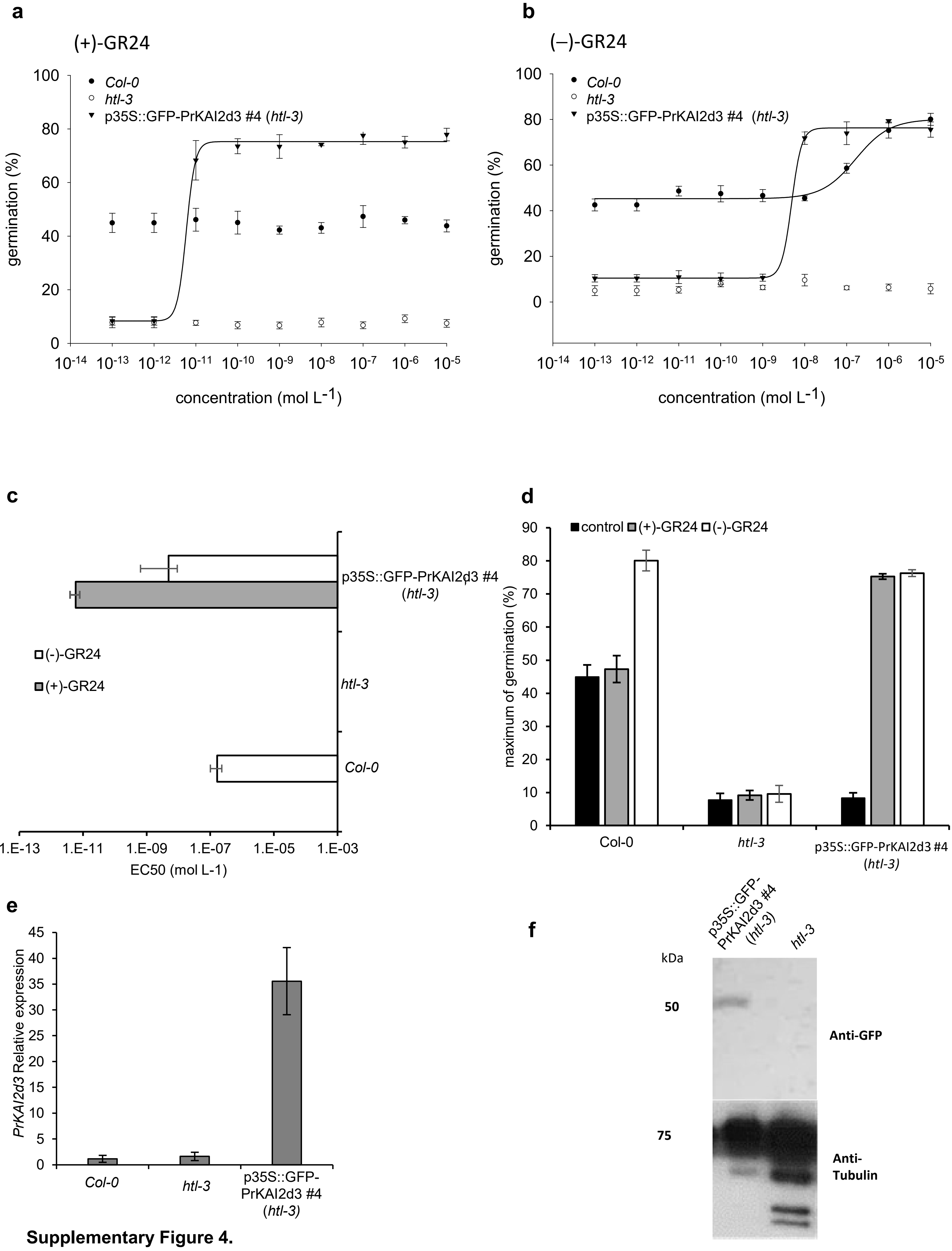

**Supplementary Figure 4.**

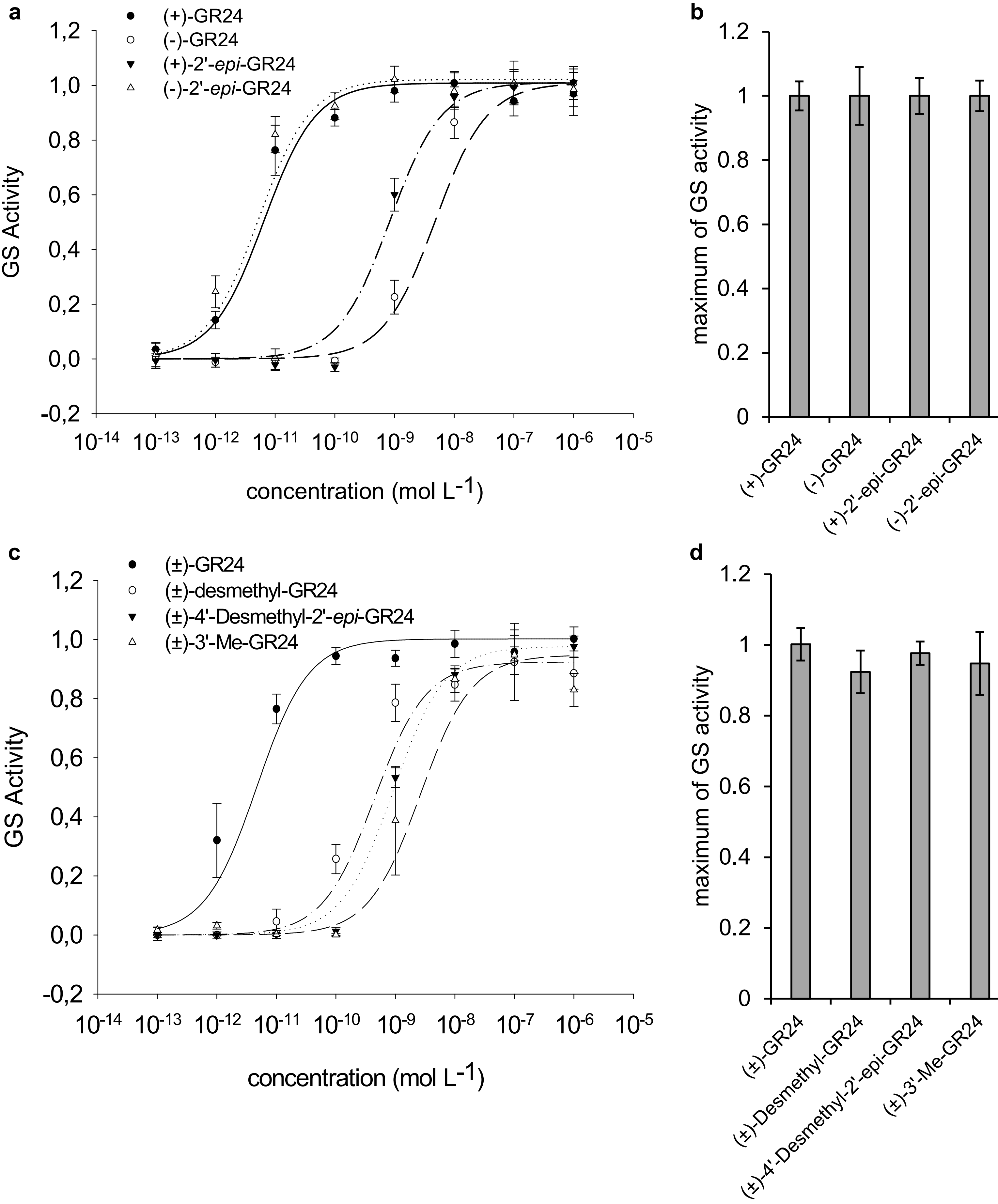

**Supplementary Figure 5.**

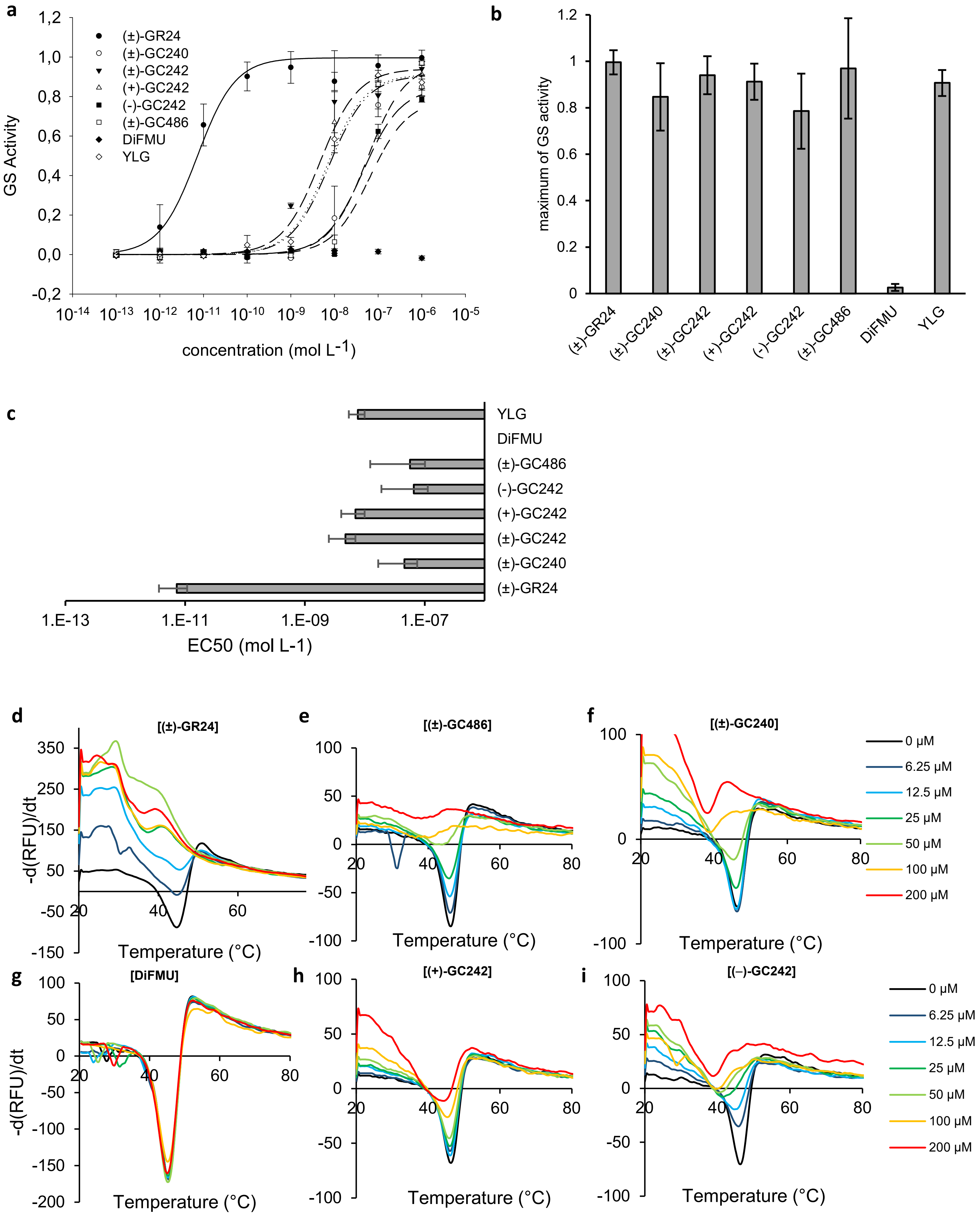

**Supplementary Figure 6.**

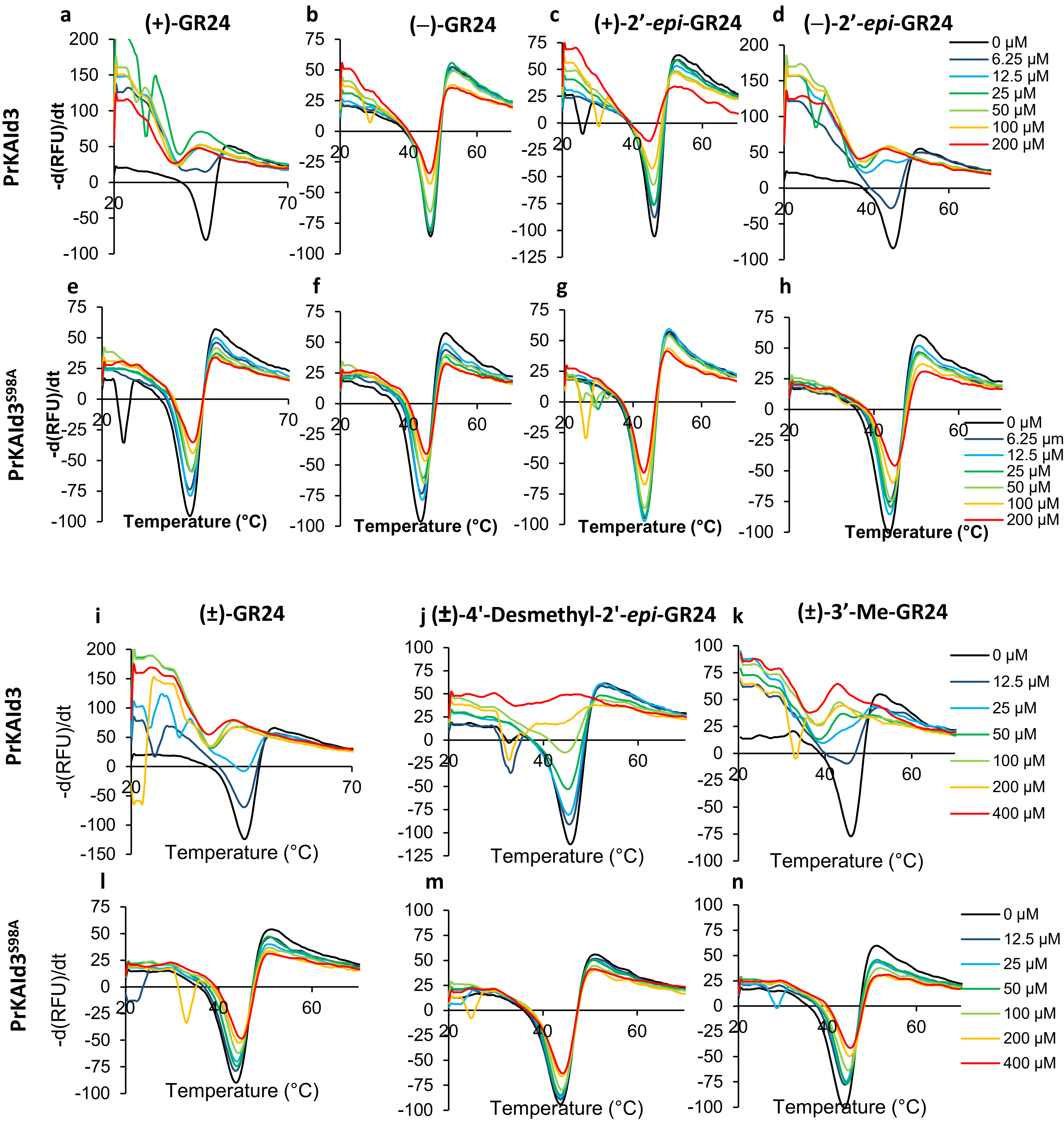

**Supplementary Figure 7.**

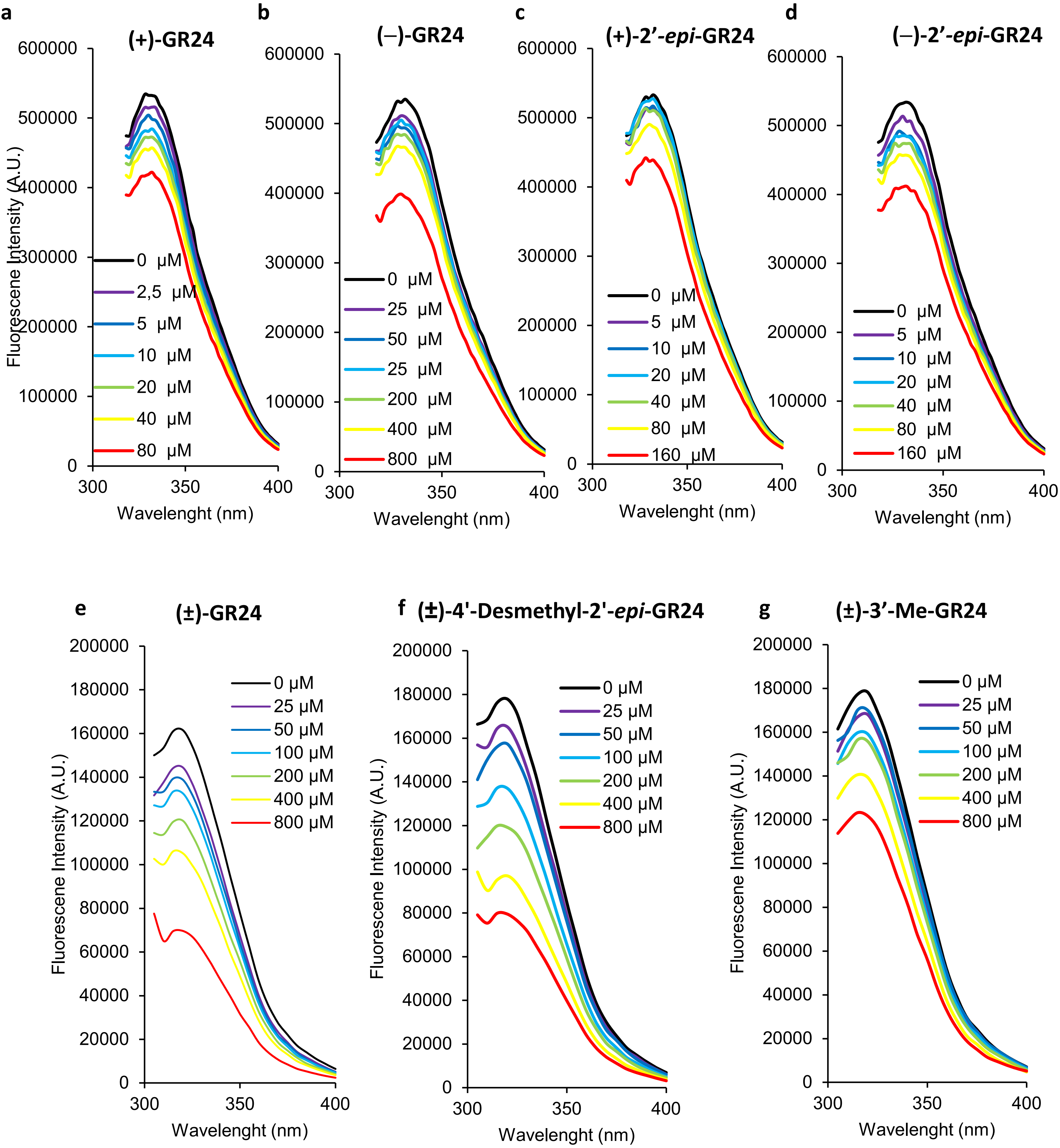

**Supplementary Figure 8.**

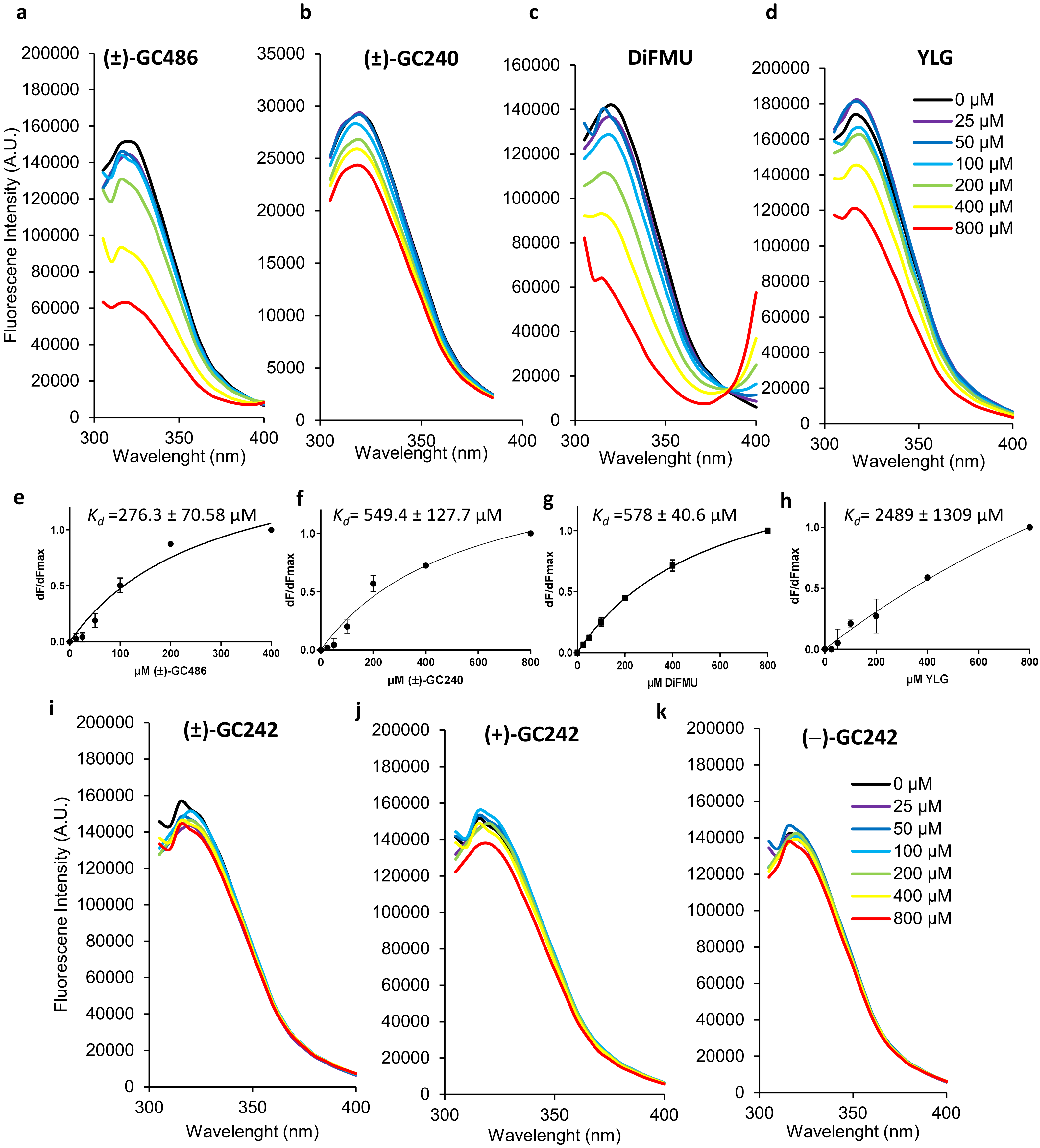

**Supplementary Figure 9.**

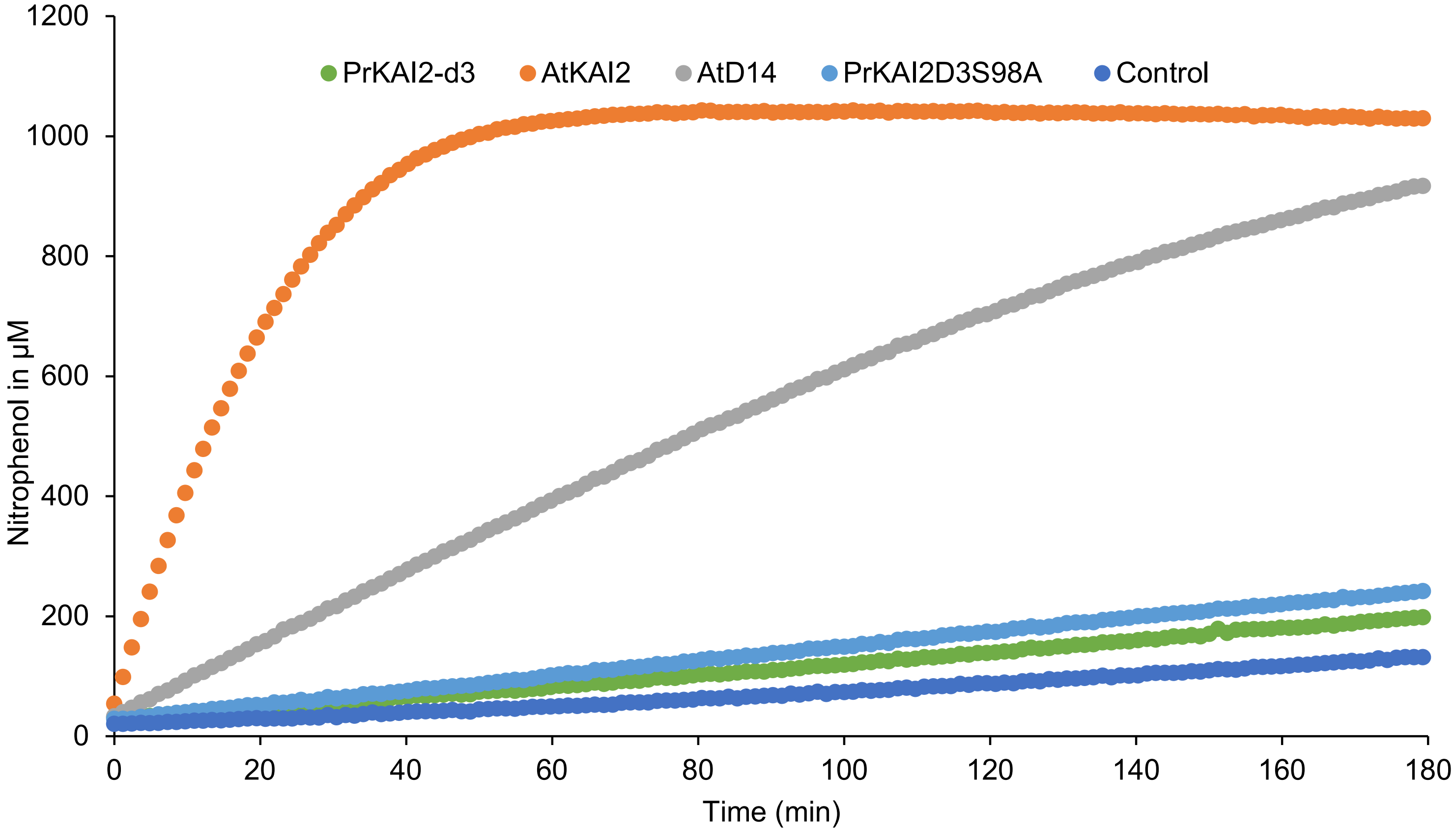

**Supplementary Figure 10.**

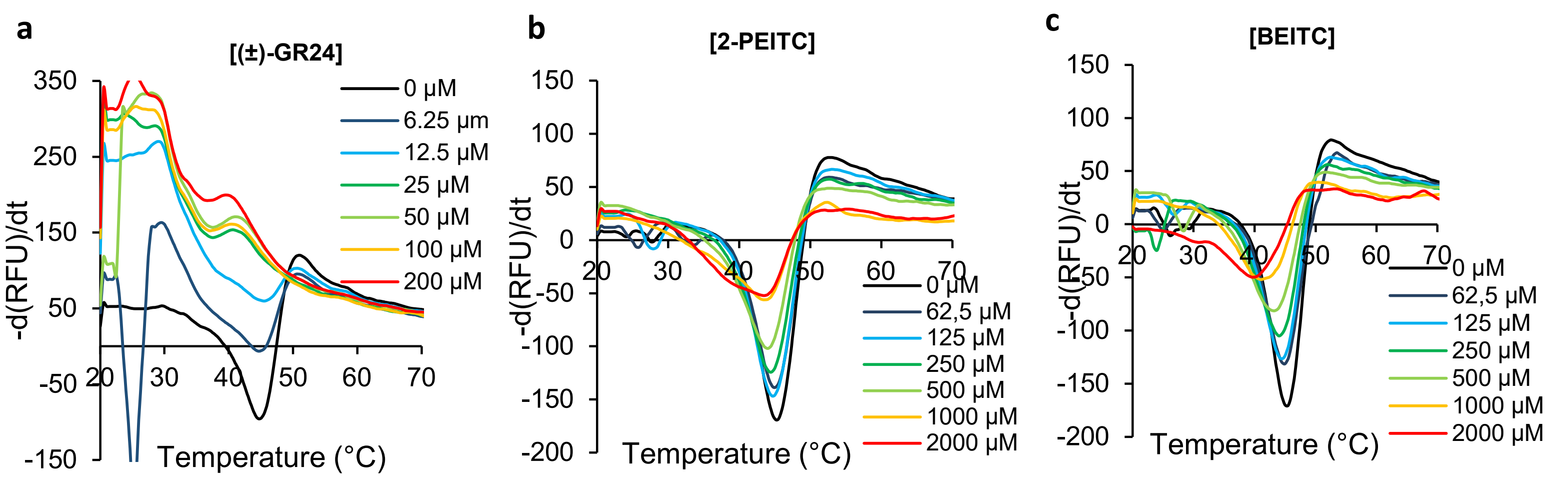

**Supplementary Figure 11.**

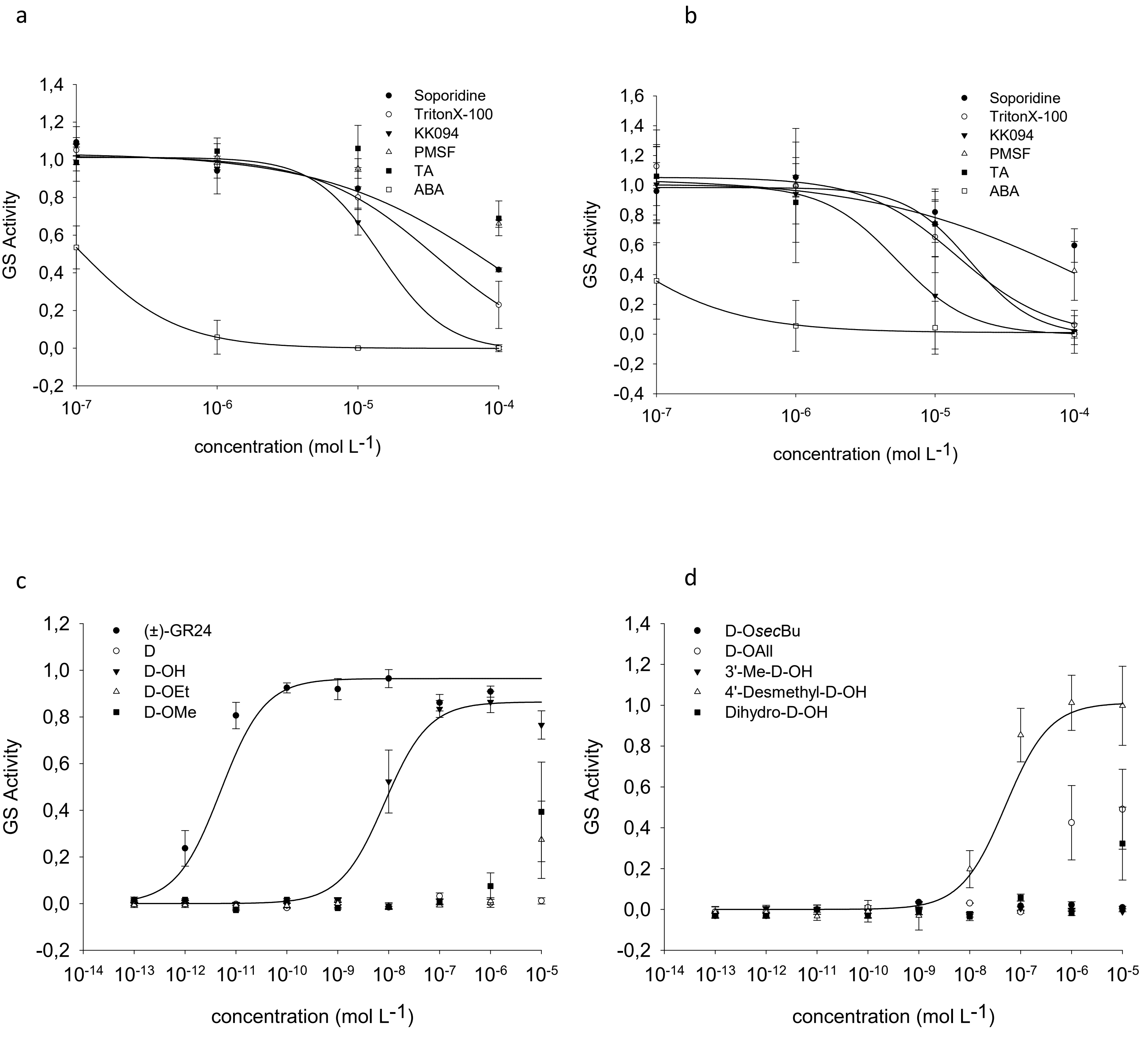

**Supplementary Figure 12.**

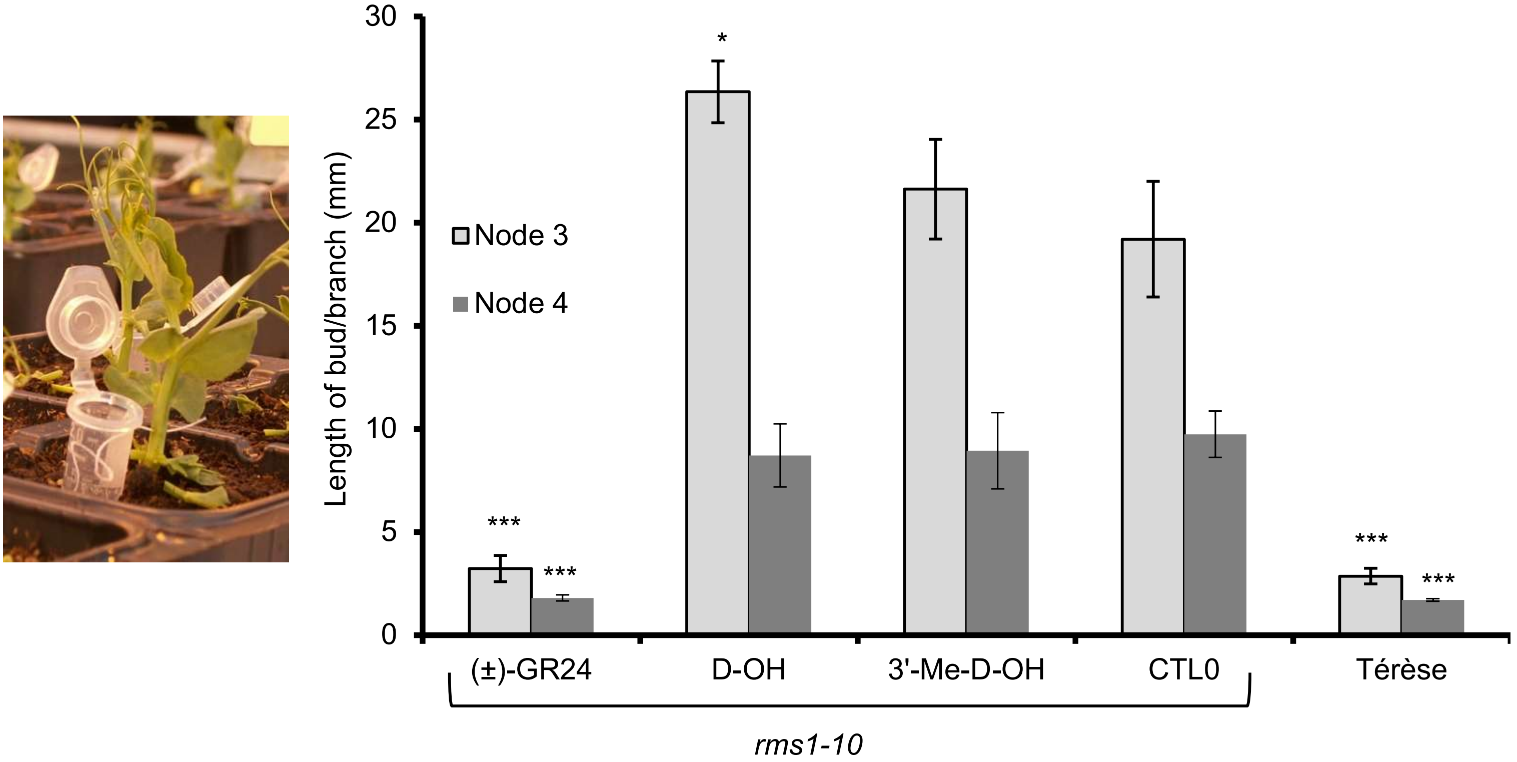

**Supplementary Figure 13.**

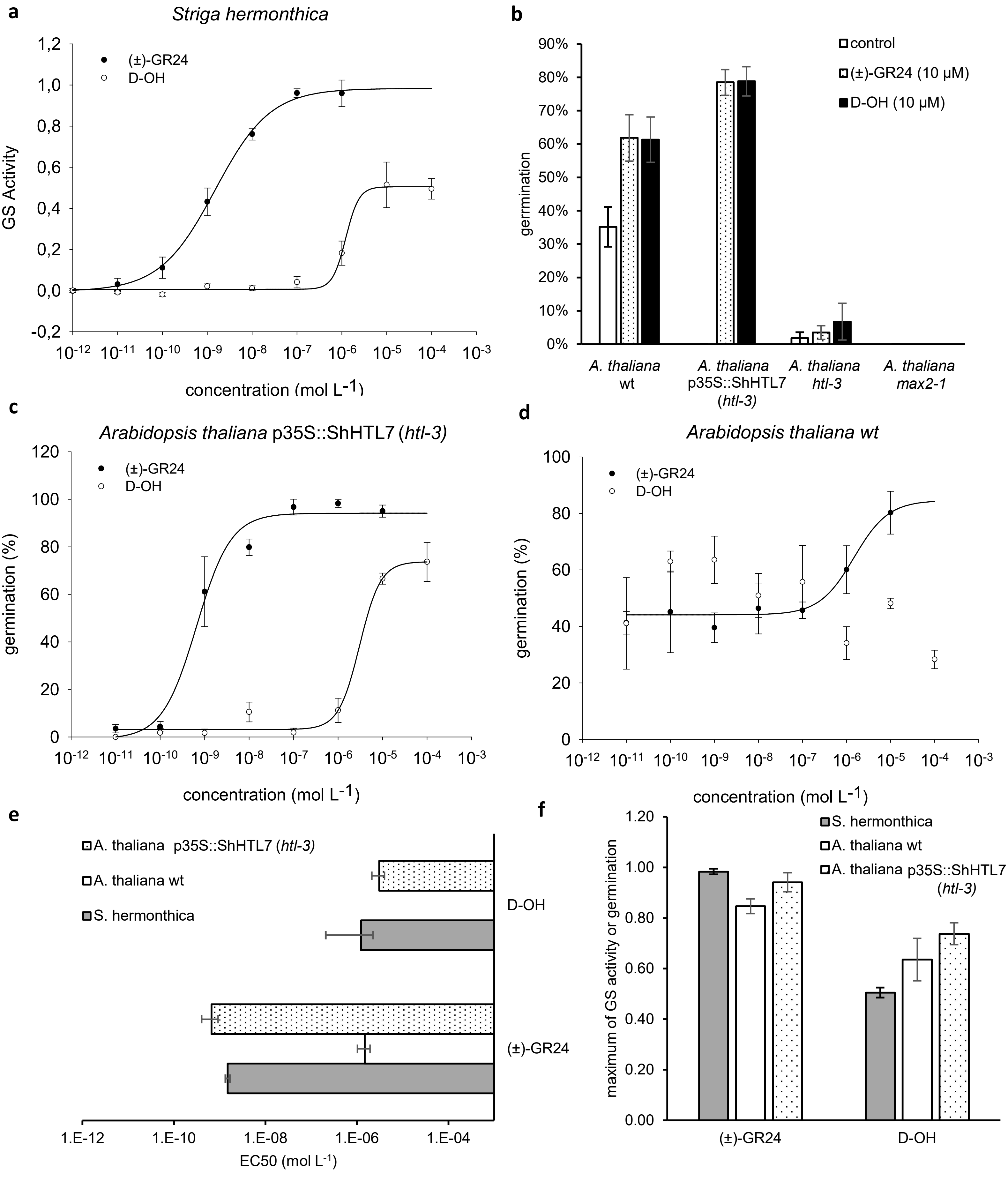

**Supplementary Figure 14.**

**Supplementary Table 1.**

| species | compound | EC50 (M) | SE | %max | SE |
|---|---|---|---|---|---|
| *A. thaliana* Col-0 | (+)-GR24 | nd | | 47% | 4% |
| *A. thaliana* Col-0 | (-)-GR24 | 1.6494E-07 | 6.4078E-08 | 80% | 3% |
| *A. thaliana htl-3* | (+)-GR24 | nd | | 9% | 1% |
| *A. thaliana htl-3* | (-)-GR24 | nd | | 10% | 3% |
| *A. Thaliana* p35S::GFP-PrKAI2d3 (*htl-3*) | (+)-GR24 | 5.909E-12 | 1.975E-11 | 75% | 1% |
| *A. thaliana* p35S::GFP-PrKAI2d3 (*htl-3*) | (-)-GR24 | 4.8909E-09 | 4.2497E-09 | 76% | 1% |
| *Striga hermonthica* | (±)-GR24 | 1.5E-09 | 1.6E-10 | 98% | 1% |
| *Striga hermonthica* | D-OH | 1.2E-06 | 1.0E-06 | 51% | 2% |
| *A. thaliana* Col-0 | (±)-GR24 | 1.5E-06 | 4.6E-07 | 85% | 3% |
| *A. thaliana* Col-0 | D-OH | nd | | 64% | 8% |
| *A. thaliana* p35S::ShHTL7 (*htl-3*) | (±)-GR24 | 6.6E-10 | 2.6E-10 | 94% | 4% |
| *A. thaliana* p35S::ShHTL7 (*htl-3*) | D-OH | 3.1E-06 | 9.4E-07 | 74% | 4% |

### Supplementary Table 2.

| compound | EC50 (M) | SE | %max | SE |
|---|---|---|---|---|
| (+)-GR24 | 6.5E-12 | 3.0E-12 | 100% | 5% |
| (-)-GR24 | 4.8E-09 | 2.3E-09 | 100% | 9% |
| (+)-2'-*epi*-GR24 | 8.4E-10 | 2.8E-10 | 100% | 6% |
| (-)-2'-*epi*-GR24 | 5.3E-12 | 2.6E-12 | 100% | 5% |
| (±)-GR24 | 4.7E-12 | 2.6E-12 | 100% | 5% |
| (±)-Desmethyl-GR24 | 4.5E-10 | 2.0E-10 | 92% | 6% |
| (±)-4'-Desmethyl-2'-*epi*-GR24 | 9.2E-10 | 1.8E-10 | 98% | 3% |
| (±)-3'-Me-GR24 | 2.6E-09 | 1.4E-09 | 95% | 9% |
| (±)-GR24 | 7.3E-12 | 3.6E-12 | 100% | 5% |
| (±)-GC240 | 4.6E-08 | 2.9E-08 | 85% | 15% |
| (±)-GC242 | 4.7E-09 | 2.2E-09 | 94% | 8% |
| (+)-GC242 | 7.0E-09 | 2.9E-09 | 91% | 8% |
| (-)-GC242 | 6.6E-08 | 4.7E-08 | 79% | 16% |
| (±)-GC486 | 5.7E-08 | 4.4E-08 | 97% | 22% |
| DiFMU | nd | nd | 3% | 2% |
| YLG | 7.7E-09 | 2.3E-09 | 91% | 6% |
| (±)-GR24 | 5.2822E-12 | 3.432E-12 | 100% | 5% |
| 2-PEITC | 3.2448E-08 | 1.9703E-08 | 102% | 15% |
| BITC | 7.1201E-08 | 2.745E-08 | 94% | 11% |
| (±)-GR24 | 5.003E-12 | 3.1496E-12 | 100% | 5% |
| D-OH | 8.3335E-09 | 2.5891E-09 | 86% | 5% |
| D-OEt | nd | | 27% | 14% |
| D-OMe | nd | | 39% | 21% |
| D-OsecBu | nd | | 1% | 1% |
| D-OAll | nd | | 49% | 20% |
| 3'-Me-D-OH | nd | | 0% | 1% |
| 4'-Desmethyl-D-OH | 5.0054E-08 | 2.0556E-08 | 101% | 8% |
| Dihydro-D-OEt | nd | | 32% | 18% |

### Supplementary Table 3.

| | GR24 (10 nM) | | | | 2-PEITC (100 nM) | | | |
|---|---|---|---|---|---|---|---|---|
| compound | IC50(M) | SE | max. inh. | SE | IC50(M) | SE | max. inh. | SE |
| Soporidine | 1.0E-04 | 8.5E-05 | 58% | 1% | nd | | 40% | 5% |
| TritonX-100 | 3.5E-05 | 6.0E-06 | 77% | 13% | 1.5E-05 | 4.1E-06 | 94% | 6% |
| KK094 | 1.4E-05 | 4.5E-06 | 100% | 2% | 5.3E-06 | 1.3E-06 | 100% | 3% |
| PMSF | nd | | 34% | 2% | 1.0E-04 | 9.0E-05 | 57% | 20% |
| TA | nd | | 31% | 9% | 1.8E-05 | 9.5E-06 | 98% | 14% |
| ABA | 1.0E-07 | 9.4E-09 | 100% | 1% | 3.4E-08 | 2.1E-08 | 100% | 7% |

**Supplementary Table 4.**

| Primer name | purpose | Sequence (5'—3') | note |
|---|---|---|---|
| Cloning for protein expression | | | |
| PrKAI2d3_attb1_HRV3C | PrKAI2d3 coding sequence | ggggacaagtttgtacaaaaaagcaggct*ccctggaagtgctgtttcagggcccg*ATGAACATTAACAGAGACAT | Gene specifics sequences are capitalized<br>Start and Stop codons are highlighted in bold<br>Gateway recombination sites are underlined<br>protease sites are in italic |
| PrKAI2d3_attb2 | PrKAI2d3 coding sequence | ggggaccactttgtacaagaaagctgggtctcaTCAATTAGCATCTGC | |
| Cloning for complementation assay | | | |
| PrKAI2d3_attb2-F | | ggggacagctttcttgtacaaagtggcaATGAACATTAACAGAGACATCGG | |
| PrKAI2d3_attb3-R | | ggggacaactttgtataataaagttggATTAGCATCTGCAATATCATG | |
| PCR-Based mutagenesis | | | |
| PrKAI2d3_S98A-F | | ctacgtcggccacgctctgtccgccat | |
| PrKAI2d3_S98A-R | | atggcggacagagcgtggccgacgtag | |
| Q-PCR primer | | | |
| PrEF1-a-F | | ttgccgtgaaggatctgaaac | |
| PrEF1-a-R | | ccttggcagggtcgtcttta | |
| PrKAI2c-F | | ccatcacaaggccagac | |
| PrKAI2c-R | | gatttcatggcctcgaa | |
| PrKAI2d3-F | | gtggccgagtatatccatc | |
| PrKAI2d3-R | | ctgtgatctccggca | |
