## Supplementary Methods for "A *Phelipanche ramosa* KAI2 Protein Perceives enzymatically Strigolactones and Isothiocyanates"

### General experimental procedures

All non-aqueous reactions were run under an inert atmosphere (argon), by using standard techniques for manipulating air-sensitive compounds. All glassware was stored in the oven and/or was flame-dried prior to use. Anhydrous solvents were obtained by filtration through drying columns. Analytical thin-layer chromatographies (TLC) were performed on plates precoated with silica gel layers. Compounds were visualized by one or more of the following methods: (1) illumination with a short wavelength UV lamp (i.e., $\lambda$ = 254 nm), (2) spray with a $KMnO_4$ solution in $H_2O$. Flash column chromatography was performed using 40-63 mesh silica. Nuclear magnetic resonance spectra ($^1$H; $^{13}$C NMR) were recorded respectively at [500; 125] MHz on a Bruker DPX 500 spectrometer. For the $^1$H spectra, data are reported as follows: chemical shift, multiplicity (s = singlet, d = doublet, t = triplet, q = quartet, m = multiplet, bs = broad singulet, coupling constant in Hz and integration. IR spectra are reported in reciprocal centimeters (cm$^{-1}$). Mass spectra (MS) and high-resolution mass spectra (HRMS) were determined by electrospray ionization (ESI) coupled to a time-of-flight analyser (Waters LCT Premier XE).

### Preparation of GR24 isomers

(±)-2'-*epi*-GR24 and (±)-GR24 were prepared according to described procedures [1]. ((+)-GR24, (−)-GR24, (+)-2'-*epi*-GR24, (−)-2'-*epi*-GR24 were separated from (±)-2'-*epi*-GR24 and (±)-GR24 by chiral supercritical fluid chromatography as described in [2]. (±)-GR24 can be purified by semi-preparative HPLC. Semi-preparative HPLC was performed using a Interchim puriFlash® 4250 instrument, combined with a fraction collector with integrated ELSD, a PDA and a Phenomenex Luna C18, 250 × 21.2 mm, 5 µm column ($H_2O/CH_3CN$ : 6 /4) or Interchim Uptisphere Strategy SI, 250 × 21.2 mm, 5 µm column (Heptane/EtOAc : 1 /1).

### Preparations of D-OH and analogs

(±)-D-OH was prepared according to Fell & Harbridge [3]. (±)-3'-Methyl-D-OH was prepared according to Canévet & Graff [4].

### Preparation of (±)-D-OEt[5,6] by a modification of the Bayer et al. procedure[5]

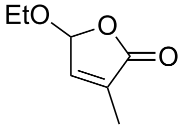


To a 0 °C solution of $TiCl_4$ (16.5 mL, 150 mmol) in anhydrous $CH_2Cl_2$ (212 mL) was added dropwise under argon a mixture of ethyl pyruvate (16.5 mL, 150 mmol) and vinyl acetate (16.5 mL, 150 mmol) in anhydrous $CH_2Cl_2$ (106 mL). When addition is complete, the suspension was further stirred for 2 hours at 0 °C. The reaction mixture was quenched with $H_2O$ (140 mL). The aqueous solution was extracted with $CH_2Cl_2$ (2 × 100 mL). The combined organic layers were washed with $H_2O$ (100 mL), brine (100 mL), dried ($Na_2SO_4$), filtered and evaporated under reduced pressure. The residue (30 g) was diluted in EtOH (345 mL), AcOH (17 mL), conc. HCl (17 mL) and stirred under reflux for 4 hours. EtOH was removed by distillation and the cooled residue was extracted with EtOAc (3 × 150 mL). The combined organic layers were washed with brine, dried ($Na_2SO_4$), filtered and evaporated under reduced pressure. The residue was distilled (120-130 °C) with Kugelrohr to give (±)-D-OEt[6] as a pale yellow liquid (9.3 g, 65.5 mmol, 44%).

**Preparation of (±)-D-OMe[7]**

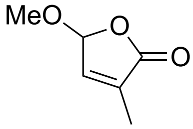


A mixture of (±)-D-O-Et (1.00 g, 7.04 mmol) and *para*-toluene sulfonic acid (250 mg, 1.31 mmol) in MeOH (44 mL) was stirred for 48 h at room temperature. MeOH was removed under reduced pressure and the residue was diluted with EtOAc (50 mL). The organic solution was washed with a $NaHCO_3$ saturated aqueous solution, brine, dried ($MgSO_4$), filtered and evaporated under reduced pressure. The residue was distilled (1 mm Hg, 70-110 °C) with Kugelrohr to give (±)-D-OMe[7] as a colourless liquid (0.81 g, 6.33 mmol, 90%).

**Preparation of (±)-Dihydro-D-OEt**

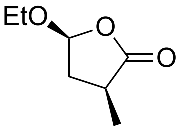


A solution of (±)-D-OEt[6] (429 mg, 3.02 mmol) and 10% Pd/C (170 mg) in EtOAc (45 mL) was stirred under $H_2$ atmosphere for 75 min at room temperature, flushed with argon and filtered on Celite. The resultant solution was evaporated under reduced pressure and distilled (1 mm Hg, 80-90 °C) with Kugelrohr to give (±)-Dihydro-D-OEt[8] as a colourless liquid (266 mg, 1.84 mmol, 61%).

**Preparation of (±)-Dihydro-D-O*sec*-Bu**

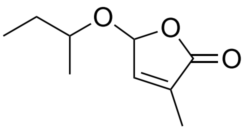


A mixture of (±)-D-O-Et (2.02 g, 14.02 mmol) and *para*-toluene sulfonic acid (503 mg) in *sec*-BuOH (40 mL) was stirred for 3 h at reflux. *sec*-BuOH was removed under reduced pressure and the residue was diluted with EtOAc (200 mL). The organic solution was washed with a 5% $NaHCO_3$ aqueous solution (2 × 50 mL), dried ($MgSO_4$), filtered and evaporated under reduced pressure. The residue was distilled (1 mm Hg, 70-110 °C) with Kugelrohr to give (±)-D-O*sec*-Bu as a colourless liquid (1.11 g, 6.52 mmol, 46%). Mixture of 2 diastereomers (1:1). $^1$H NMR ($CDCl_3$, 300 MHz): $\delta$ (ppm) 0.87 (t, $J$ = 7.5 Hz, 3H), 0.90 (t, $J$ = 7.5 Hz, 3H), 1.19 (d, $J$ = 6.0 Hz, 3H), 1.23 (d, $J$ = 6.0 Hz, 3H), 1.44-1.64 (m, 4H), 1.89-1.91 (m, 6H), 3.75-3.85 (m, 2H), 5.81-5.83 (m, 1H), 5.83-5.86 (m, 1H), 6.73-6.76 (m, 2H). $^{13}$C NMR ($CDCl_3$, 75 MHz): $\delta$ (ppm) 9.7 ($CH_3$), 9.8 ($CH_3$), 10.72 ($CH_3$), 10.76 ($CH_3$), 19.4 ($CH_3$), 20.9 ($CH_3$), 29.5 ($CH_2$), 29.9 ($CH_2$), 77.5 (CH), 77.9 (CH), 100.2 (CH), 101.7 (CH), 133.9 (2C), 143.3 (CH), 143.5 (CH), 172.28 (C), 172.30 (C). HRMS (ESI): Calculated for $C_9H_{15}O_3$ $[M + H]^+$: 171.1021. Found: 171.1019.

**Preparation of (±)-Dihydro-D-OAll**

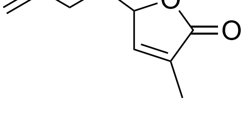


A mixture of (±)-D-O-Et (1.01 g, 7.13 mmol) and *para*-toluene sulfonic acid (250 mg, 1.31 mmol) in allylic alcohol (40 mL) was stirred for 24 h at room temperature. Allylic alcohol was removed under reduced pressure and the residue was diluted with EtOAc (100 mL). The organic solution was washed with a 5% $NaHCO_3$ aqueous solution (2 × 50 mL), dried ($MgSO_4$), filtered and evaporated under reduced pressure. The residue was distilled (20 mm Hg, 90 °C) with Kugelrohr to give (±)-D-OAll as a colourless liquid (719 mg, 4.66 mmol, 65%). $^1$H NMR ($CDCl_3$, 300 MHz): $\delta$ (ppm) 1.90 (s, 3H), 4.16 (dd, $J$ = 12.5, 6.5 Hz, 1H), 4.32 (dd, $J$ = 12.5, 5.5 Hz, 1H), 5.22 (d, $J$ = 10.5 Hz, 1H), 5.30 (dd, $J$ = 17.0, 3.0 Hz,

1H), 5.79-5.81 (bs, 1H), 5.82-5.95 (m, 1H), 6.78-6.79 (bs, 1H). $^{13}C$ NMR ($CDCl_3$, 75 MHz): $\delta$ (ppm) 10.7 ($CH_3$), 70.8 ($CH_2$), 100.7 (CH), 119.0 ($CH_2$), 133.0 (CH), 134.3 (C), 143.0 (CH), 172.0 (C). IR (film): $\nu$ ($cm^{-1}$) 3085, 2983, 2928, 1764, 1668, 1649. HRMS (ESI): Calculated for $C_8H_{11}O_3$ $[M + H]^+$: 155.0708. Found: 155.0703.

**Preparation of (±)-3'-desmethyl-D-OH**

To a solution of (±)-3'-desmethyl-D-Br[9] (200 mg, 1.23 mmol) was added an aqueous solution of KOH (2 mL, 2 M). The resultant mixture was stirred at room temperature for 2 h, neutralized with an aqueous solution of HCl (1 M) and extracted with EtOAc (3 × 15 mL). The combined organic layers were dried ($Na_2SO_4$), filtered and evaporated under reduced pressure. The residue was chromatographied on silica gel (Heptane/ EtOAc 100:0 to 1:1) to give (±)-3'-desmethyl-D-OH[10] as a white solid (45 mg, 37%).


1 Mangnus, E. M., Dommerholt, F. J., Dejong, R. L. P. & Zwanenburg, B. Improved Synthesis of Strigol Analog GR24 and Evaluation of the Biological-Activity of Its Diastereomers. *J. Agric. Food. Chem.* **40**, 1230-1235, doi:10.1021/jf00019a031 (1992).

2 de Saint Germain, A., Clavé, G., Badet-Denisot, M.-A., Pillot, J.-P., Cornu, D., Le Caer, J.-P., Burger, M., Pelissier, F., Retailleau, P., Turnbull, C., Bonhomme, S., Chory, J., Rameau, C. & Boyer, F.-D. An histidine covalent receptor and butenolide complex mediates strigolactone perception. *Nat. Chem. Biol.* **12**, 787-794, doi:10.1038/nchembio.2147 (2016).

3 Fell, S. C. M. & Harbridge, J. B. 2,5-dimethoxy-2,5-dihydrofuran: A convenient synthon for a novel mono-protected glyoxal; synthesis of 4-hydroxybutenolides. *Tetrahedron Lett.* **31**, 4227-4228, doi:10.1016/S0040-4039(00)97588-9 (1990).

4 Canévet, J. C. & Graff, Y. Réactions de friedel-crafts de dérivés aromatiques sur des composés dicarbonylés-1,4éthyléniques-2,3.ii alkylations par quelques hydroxy-5 ou chloro-5 dihydro-2,5 furannones-2. nouvelle méthode de synthèse des acides 1h-indènecarboxyliques-1. *Tetrahedron* **34**, 1935-1942, doi:10.1016/0040-4020(78)80100-8 (1978).

5 Bayer, T. S., Davidson, E. A. & Hleba, Y. Methods for hydraulic enhancement of crops. PCT/US2016/029080 (2016).

6 Wigchert, S. C. M. & Zwanenburg, B. A critical account on the reception of Striga seed germination. *J. Agric. Food. Chem.* **47**, 1320-1325, doi:10.1021/jf980926e (1999).

7 Chapleo, C. B., Svanholt, K. L., Martin, R. & Dreiding, A. S. Synthesis of Bromo-substituted 2-Buten- and 2-Penten-4-olides. *Helv. Chim. Acta* **59**, 100-107, doi:10.1002/hlca.19760590109 (1976).

8 Redon, S., Pannecoucke, X., Franck, X. & Outurquin, F. Synthesis and oxidative rearrangement of selenenylated dihydropyrans. *Org. Biomol. Chem.* **6**, 1260-1267, doi:10.1039/b718825k (2008).

9 Wolff, S. & Hoffmann, H. M. R. Aflatoxins revisited - convergent synthesis of the ABC-moiety. *Synthesis*, 760-763, doi:10.1055/s-1988-27700 (1988).

10 Byun, J., Huang, W., Wang, D., Li, R. & Zhang, K. A. I. CO2 -Triggered Switchable Hydrophilicity of a Heterogeneous Conjugated Polymer Photocatalyst for Enhanced Catalytic Activity in Water. *Angew. Chem. Int. Ed.* **57**, 2967-2971, doi:10.1002/anie.201711773 (2018).
